# Structural basis for CDC42 and RAC activation by the dual specificity GEF DOCK10

**DOI:** 10.1101/2022.06.15.496229

**Authors:** Danni Fan, Jing Yang, Nora Cronin, David Barford

**Affiliations:** Division of Structural Biology, Institute of Cancer Research, London, SW3 6JB, UK

**Author notes:** MRC Laboratory of Molecular Biology, Cambridge, UK.

**Keywords:** GEF, DOCK proteins, DOCK10, GTPases, RAC, CDC42, protein crystallography

## Abstract

Dedicator of cytokinesis 10 (DOCK10) is a guanine nucleotide exchange factor (GEF) belonging to the DOCK family that activate RHO GTPases. DOCK10 controls amoeboid migration and IL-4 induced B cell activation. Recent structural studies of the catalytic DHR2 domains of DOCK-family GEFs DOCK2 and DOCK9 in the presence of nucleotide-free RAC and CDC42 respectively, have revealed a conserved mechanism of nucleotide exchange distinct from the large family of Dbl homology (DH) GEFs. Less is known from a biochemical and structural perspective, about DOCK10. We have determined the crystal structure of the apo-state of the DHR2 domain of DOCK10 (DOCK10^DHR2^), which provides the first structural model for the apo-state of a DHR2 domain for the entire DOCK family. It demonstrates the conformational changes within DOCK10^DHR2^ on engaging the GTPase, and associated induced conformational changes of the GTPase required to stimulate nucleotide release. We also report on the unexpected dual specificity of DOCK10^DHR2^ which directly interacts with RAC and CDC42 and induces the nucleotide exchange of both *in vitro*. This dual specificity is unique to DOCK10 within its evolutionarily related D subfamily. Structural studies of DOCK10^DHR2^ in complex with RAC revealed an intriguing 2:1 stoichiometry between DOCK10^DHR2^ and the GTPase, that differs from the canonical 1:1 stoichiometry for all other known DOCK-GTPase complexes. This was confirmed in solution using a variety of biophysical techniques. The identification of a novel mode of interaction between DOCK10^DHR2^ and the RAC GTPase provides new insights into how DOCK family GEFs discriminate between CDC42, RAC and RHOA.

---

Dedicator of cytokinesis (DOCK) family guanine nucleotide exchange factors (GEFs) are specific activators of RHO GTPases through catalyzing the exchange of GDP for GTP. In mammals, the family comprises eleven members organized into four subfamilies on the basis of sequence relatedness, domain structure and specificity for either RAC or CDC42 (1,2). DOCK10, together with DOCK9 and DOCK11, belongs to the DOCK-D subfamily, which is the only DOCK family comprising an N-terminal pleckstrin homology (PH) domain (2-6). DOCK-D members have varied tissue distributions. Unlike DOCK9, DOCK10 and DOCK11 are both predominantly expressed in hematopoietic tissues, particular in peripheral blood leukocytes. DOCK10 is highly expressed in lymph nodes and is also expressed in some non-hematopoietic tissues such as brain and kidney (6-8).

DOCK10 plays crucial functions in amoeboid migration and in regulating the epithelial to mesenchymal transition (EMT). Depleting DOCK10 shifted the migration mode from amoeboid to mesenchymal (5). A separate study showed that DOCK10 is over-expressed in aggressive poorly differentiated papillary thyroid carcinomas. The extensive local invasion or synchronous remote metastases are consistent with the ability of DOCK10 to trigger amoeboid movement (9). In addition, it was found that IL-4 consistently increased DOCK10 levels significantly in the cytoplasm and nucleus of chronic lymphoid leukemia (CLL) and normal peripheral blood B cells, but not in T cells. In contrast, IL-4 did not affect the protein level of DOCK9 or DOCK11 in CLL cells (8). This suggests a role for DOCK10 in IL-4 induced B cell activation, which links IL-4 signaling and the RHO GTPase function in B cells. DOCK10 induces ruffles and filopodia when expressed in HeLa cells (10,11), implicating a role for DOCK10 in controlling cell migration and consistent with findings that DOCK10 may play a role in cell invasion in breast cancer (12).

DOCK10, in common with other DOCK-D subfamily GEFs, comprises three conserved domains: Pleckstrin Homology (PH), DOCK Homology Region 1 (DHR1) and DOCK Homology Region 2 (DHR2) domains. The DHR1 and PH domains have been implicated in membrane localization and allosteric modulation of exchange activity through phosphoinositide binding (13,14). The DHR2 domain of DOCK10 is the catalytic domain, which was assumed to specifically activate CDC42 (3). Recent structural studies of other DOCK family GEFs have revealed a conserved catalytic mechanism of guanine nucleotide exchange in RHO GTPases. Crystal structures of DOCK9^DHR2^-CDC42 and DOCK2^DHR2^-RAC1 complexes revealed that conserved α-helices of the DHR2 domain engage and reshape the switch 1 region of their cognate GTPases. The altered conformation of the GTPase disrupts both magnesium and GDP binding (15-17). These studies also shed light on how DHR2 domains couple to different nucleotide bound states of GTPases. However, some important questions have remained unanswered regarding the structural basis of specificity of these GEFs for their GTPase substrates.

In this study we address the structure-function relationships of the DHR2 domain of DOCK10. The crystal structure of apo-state of human DOCK10^DHR2^ was determined, which provides the first structural model for the apo-state of the DHR2 domain for the entire DOCK family. It demonstrates that in the unliganded apo-state (without GTPase), the DOCK10^DHR2^ domain undergoes significant restructuring. Furthermore, a novel interaction between DOCK10 and the RAC GTPase has been identified. To our knowledge, this is the first example of a DOCK protein from the DOCK-D subfamily that has been confirmed *in vitro* to have dual specificity, catalyzing nucleotide exchange of both CDC42 and RAC1/3. Previously DOCK7 was assigned to have dual specificity for CDC42 and RAC *in vivo* (18,19), and recently *in vitro* (20), although with very low rate enhancements of three-fold for CDC42 and two-fold for RAC1.

## Results

### Structural and functional analysis of human apo-DOCK10^DHR2^

The crystal structure of human apo-DOCK10^DHR2^ was determined at 5 Å resolution, using molecular replacement and DEN refinement (26) (**Fig. 1A, B**). The structure of the DHR2 domain is organized into three lobes of roughly equal size (lobes A, B, and C on the basis of previous nomenclature (15)). Lobe A consists of an antiparallel array of five α-helices. The dimer interface of DOCK10^DHR2^ is generated by the α4 and α5 helices of lobe A, whereas lobes B and C combine to generate the binding interface for the GTPase. The apo-state structure of DOCK10^DHR2^ was investigated using SAXS (**Fig. 1C, D**). In **Fig. 1D** the theoretical SAXS curve, calculated from the atomic structure by CRYSOL (28), is compared with the experimental SAXS data of the apo DOCK10^DHR2^ domain. The two curves match closely. A chi value of 2.03 suggests that the crystal structure agrees closely with the experimental scattering data, and is therefore a good representation of the solution structure of the apo DOCK10^DHR2^ domain.

**Figure 1.**
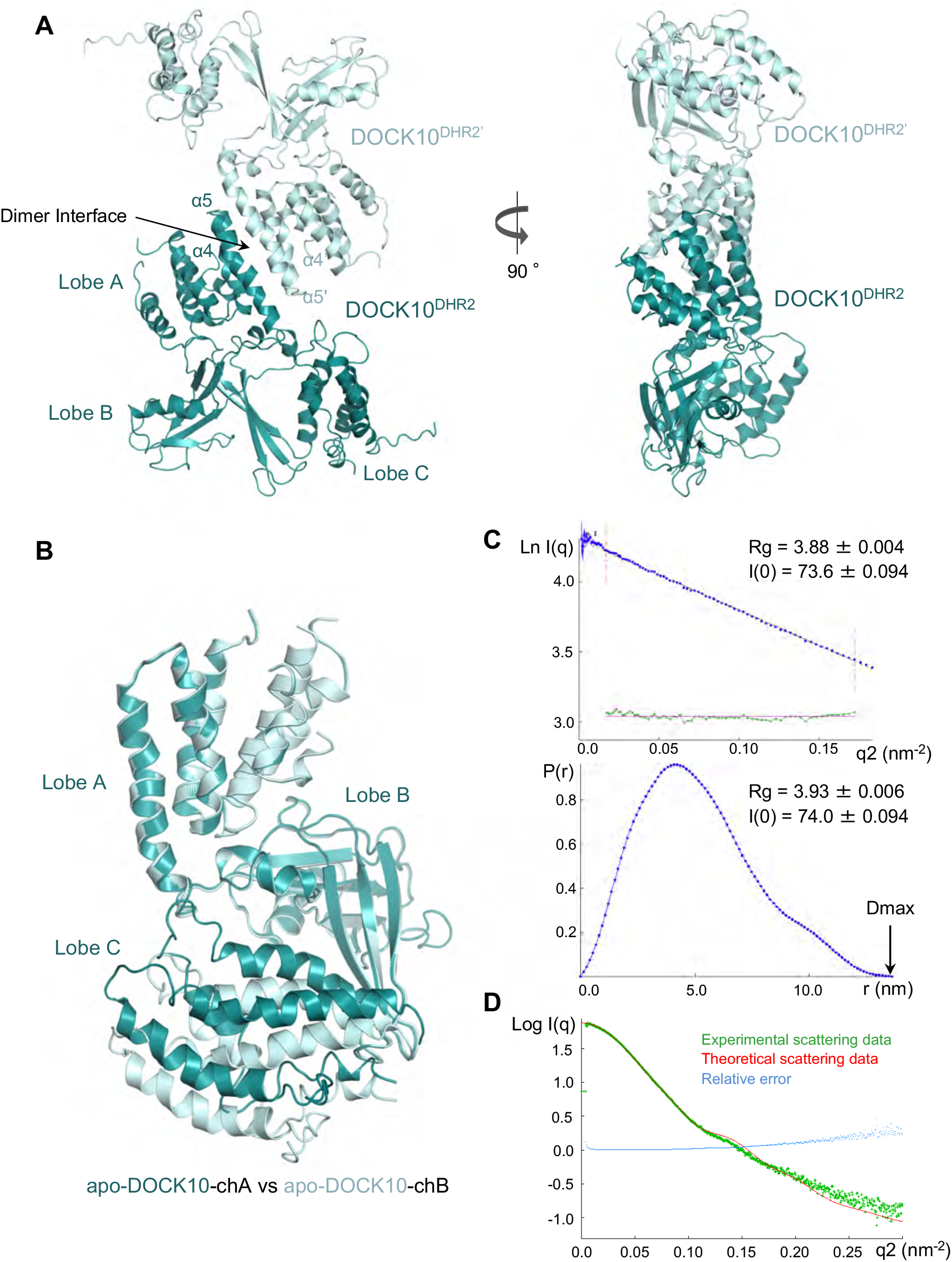
Structural analysis of apo-DOCK10^DHR2^. **A**. Overall fold of apo-DOCK10^DHR2^ dimer in the crystal structure. One molecule of DOCK10^DHR2^ is colored in cyan and the other is colored light cyan. **B**. Superimposition of two subunits of apo-DOCK10^DHR2^ present in the DOCK10^DHR2^ dimer. The two subunits were superimposed on lobes A and B. These adopt a rigid conformation, and show that the position of lobe C varies relative to lobes A-B between the two subunits of the dimer. **C**. Analysis of SAXS data. (Top) Guinier region of X-ray scattering curve of DOCK10^DHR2^. (Bottom) Distance distribution function of DOCK10^DHR2^. **D**. Theoretical SAXS profiles of DOCK10^DHR2^ (red) fixed against the experimental SAXS profile (green), generated with the program CRYSOL (28). The CRYSOL Chi value for the match of the crystal structure with the experimental data is 2.03 (value in parentheses), where Chi value around 1 is optimal.

Unlike previously determined structures of DOCK^DHR2^ in complex with their cognate GTPases (DOCK2 and DOCK9) (15,16), the structure of the apo DOCK10^DHR2^ dimer is asymmetric. This asymmetry results from conformational variations involving the orientation of lobe C relative to lobes A and B, which together form a rigid structure (**Fig. 1B**). The distinct conformations of lobe C in the absence of the bound GTPase indicate that lobe C is mobile in solution. The crystal contacts have probably enabled us to view two conformational states from possibly a larger number of states present in solution. Lobes B and C of the GTPase-binding interface are freely accessible in the apo-state of the DOCK10^DHR2^ structure. As shown later, engaging either CDC42 or RAC restricts lobe C relative to lobes A and B to essentially one conformation.

### Functional analysis of the DOCK10^DHR2^ domain in complex with its RHO GTPases

Mass-spectrometry results for the pull-down samples of Strep-tagged DOCK10^DHR2^ from HEK293 cells showed the unexpected presence of RAC3 (data not shown). To address whether the interaction was direct, and to eliminate the possibility that other eukaryotic protein(s) could mediate the interaction, purified proteins expressed in *E. coli* were further tested for binding *in vitro*. Strep-tagged DOCK10^DHR2^ was tested in pull down assays with the RHO GTPases RHOA, RAC3 and CDC42. The pull-down samples were assessed on SDS-PAGE (**Fig. 2A**). Both CDC42 and RAC3 were conclusively pulled down by DOCK10^DHR2^, suggesting direct binding. There was no detectable binding of RHOA by DOCK10^DHR2^.

**Figure 2.**
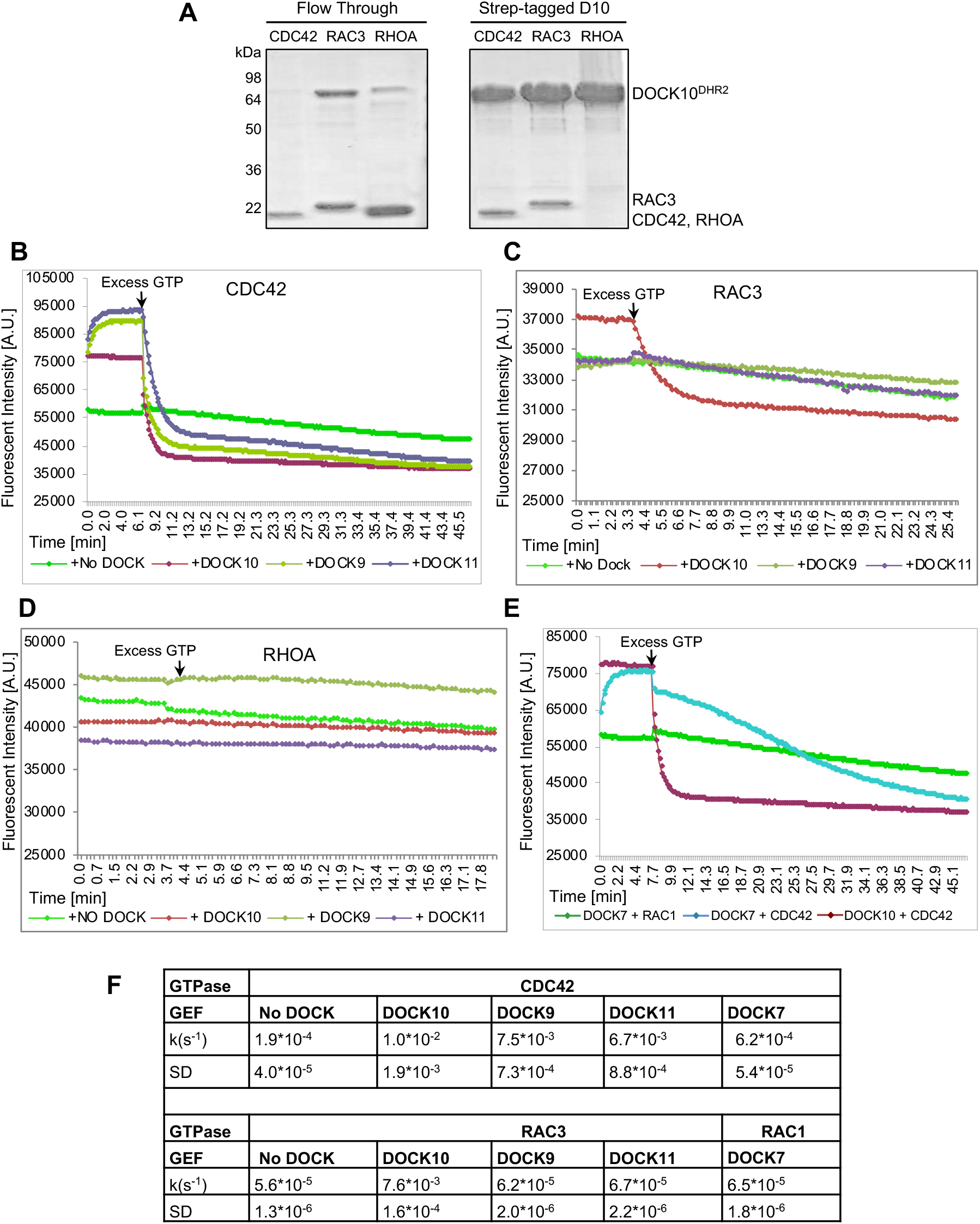
A. Protein interactions analysis. Detecting specific and stable direct binding of DOCK10^DHR2^ to GTPases in a pull-down experiment. Flow-through samples were collected; resins were washed extensively and boiled to be analyzed on reducing SDS-PAGE. CDC42, RAC3 and RHO A were detected as bands at 19.8 kDa, 21.4 kDa and 20.2 kDa respectively. **B-E**. Kinetics analysis of DOCK^DHR2^ domain catalyzed nucleotide release from GTPases. Raw data are plotted. Arrow indicates addition of excess GTP. The first three graphs shown are plots of the release of mant-GDP in the presence of different DOCK proteins, from CDC42, RAC3 or RHO A respectively. The last graph shown are the kinetic data for DOCK7^DHR2^ induced mant-GDP release from CDC42 or RAC1 in the presence of excess GTP. DOCK10 ^DHR2^ catalyzed nucleotide release from CDC42 was used as a control. **F**. Table of kinetic data for mant-GDP release from GTPases in the presence or absence of the DOCK DHR2 domain, upon addition of excess GTP. Mean and standard deviation (SD) of triplicate experiments (assuming normal distribution of data).

### Assessing the GEF activity of the other DOCK-D family members towards RHO GTPases

Previously, members of the DOCK-D subfamily were assigned as CDC42-specific GEFs. Although DOCK9 and DOCK11 were shown to be specific for CDC42 (Cote and Vuori, 2002; Lin et al., 2006; Kilkurni et al., 2011), the GTPase specificity of DOCK10 had not been previously determined. We therefore performed a comparative study of the activity of DOCK10^DHR2^ towards CDC42, RAC3 and RHO. DOCK9 and DOCK11 were included as controls.

To determine the catalytic activity of DOCK10^DHR2^ towards CDC42 and RAC3, a fluorescence-based GEF assay was performed to monitor the release of fluorescent mant-GDP upon addition of excess GTP. This showed that DOCK10^DHR2^ catalyzed nucleotide exchange activity on both CDC42 and RAC3, with rate enhancements of 53-and 136-fold, respectively (**Fig. 2B, C**). DOCK10^DHR2^ can also bind and activate RAC1 (data not shown). Thus, the capability of DOCK10^DHR2^ to catalyze nucleotide exchange reactions for CDC42, RAC3 and RAC1 correlated exactly with its ability to bind CDC42, RAC3 and RAC1 under nucleotide-free conditions. Taken together, these data suggested that DOCK10^DHR2^ itself binds to nucleotide free CDC42, RAC3 and RAC1, and can mediate nucleotide exchange on these three GTPases.

As shown in **Fig. 2B**, DOCK9^DHR2^ and DOCK11^DHR2^ clearly exhibited GEF activity for CDC42 (in agreement with Cote and Vuori, 2002; Lin et al., 2006; Kilkurni et al., 2011), but failed to demonstrate any GEF activity towards RAC3 and RHOA (**Fig. 2C, D**). The rates of the nucleotide exchange are summarized in **Fig. 2F**. These finding correlated well with the capabilities of DOCK9^DHR2^ and DOCK11^DHR2^ to bind to nucleotide-free CDC42, but not to RAC3 and RHOA in an *in vitro* pull-down experiment (data not shown). Therefore, despite the high structural similarity of the DOCK-D subfamily members, DOCK10^DHR2^ is the only DOCK-D family member that demonstrates dual specificity towards both CDC42 and RAC.

On the other hand, DOCK7^DHR2^, a DOCK-C subfamily GEF which was previously assigned to have dual specificity towards CDC42 and RAC1 *in vivo*, only catalyzed extremely slow nucleotide exchange on CDC42 (3-fold higher than the uncatalyzed rate). We did not observe catalyzed-nucleotide exchange on RAC1 (**Fig. 2E**). The three-fold enhanced rate of GEF activity towards CDC42 agrees with the recent work of Shirouzu and colleagues (20), however, unlike the latter study, which reported a two-fold rate enhancement of nucleotide exchange for RAC1, we did not detect DOCK7^DHR2^ stimulation of nucleotide exchange for RAC1. DOCK10^DHR2^ has similar activities towards CDC42 as DOCK9^DHR2^ and DOCK11^DHR2^, which are 16-fold higher than that of DOCK7^DHR2^ (**Fig. 2F**).

### Structural analysis of the DOCK10^DHR2^ domain in complex with CDC42 and CDC42-GDP

To understand the dual specificity of DOCK10^DHR2^ for CDC42 and RAC, we determined crystal structures of DOCK10^DHR2^ in complex with nucleotide-free CDC42 and RAC3. The structure of DOCK10^DHR2^ in complex with nucleotide-free CDC42 was determined at 2.5 Å resolution (**Table 1**). Most of the residues of the DOCK10^DHR2^-CDC42 complex are well defined in electron density.

**Table 1.**
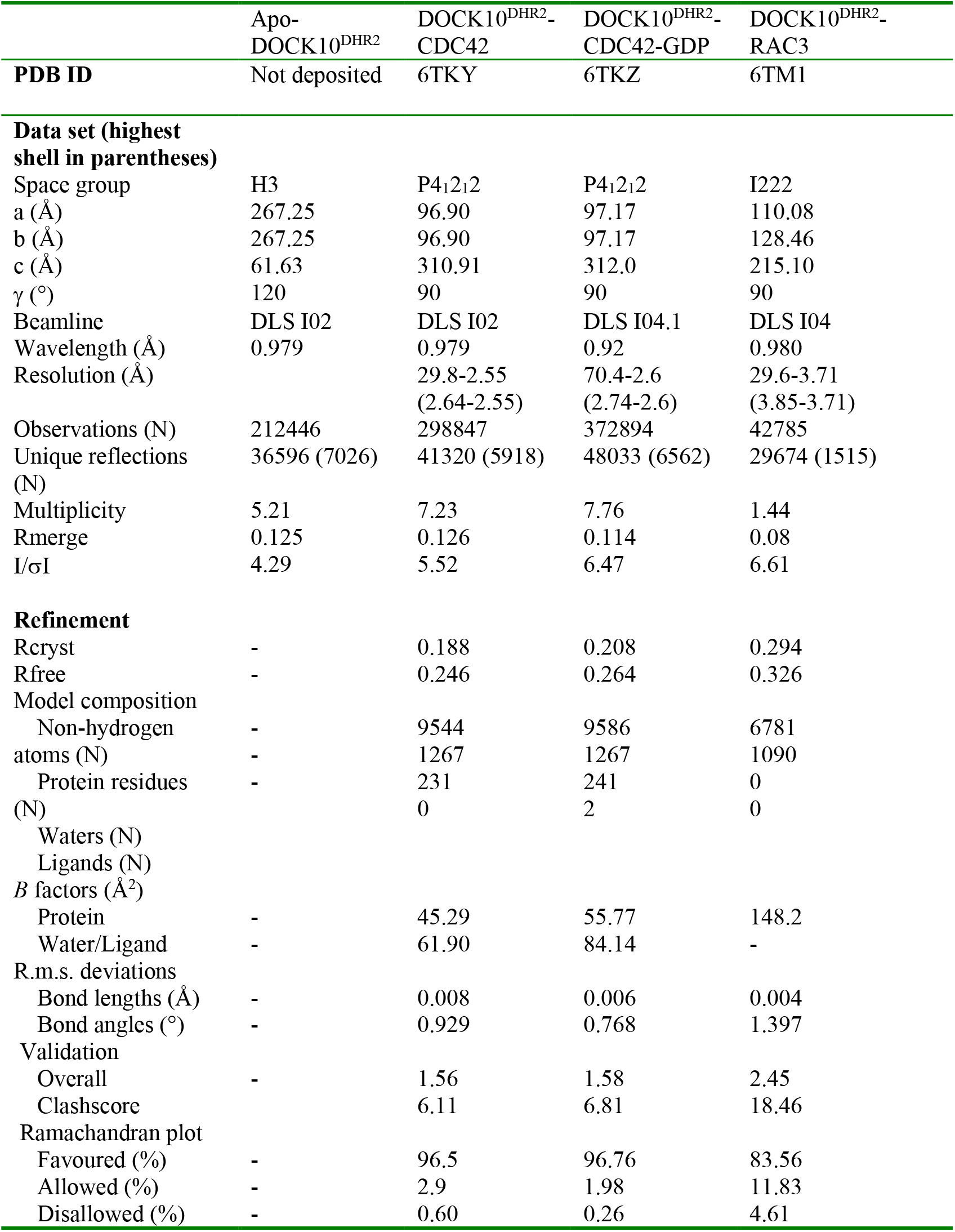
Data collection and refinement statistics.

|  | Apo-DOCK10<sup>DHR2</sup> | DOCK10<sup>DHR2</sup>-<br>CDC42 | DOCK10<sup>DHR2</sup>-<br>CDC42-GDP | DOCK10<sup>DHR2</sup>-<br>RAC3 |
| --- | --- | --- | --- | --- |
| **PDB ID** | Not deposited | 6TKY | 6TKZ | 6TM1 |
| **Data set (highest shell in parentheses)** |  |  |  |  |
| Space group | H3 | P4<sub>1</sub>2<sub>1</sub>2 | P4<sub>1</sub>2<sub>1</sub>2 | I222 |
| a (Å) | 267.25 | 96.90 | 97.17 | 110.08 |
| b (Å) | 267.25 | 96.90 | 97.17 | 128.46 |
| c (Å) | 61.63 | 310.91 | 312.0 | 215.10 |
| γ (°) | 120 | 90 | 90 | 90 |
| Beamline | DLS I02 | DLS I02 | DLS I04.1 | DLS I04 |
| Wavelength (Å) | 0.979 | 0.979 | 0.92 | 0.980 |
| Resolution (Å) |  | 29.8-2.55<br>(2.64-2.55) | 70.4-2.6<br>(2.74-2.6) | 29.6-3.71<br>(3.85-3.71) |
| Observations (N) | 212446 | 298847 | 372894 | 42785 |
| Unique reflections (N) | 36596 (7026) | 41320 (5918) | 48033 (6562) | 29674 (1515) |
| Multiplicity | 5.21 | 7.23 | 7.76 | 1.44 |
| Rmerge | 0.125 | 0.126 | 0.114 | 0.08 |
| I/σI | 4.29 | 5.52 | 6.47 | 6.61 |
| **Refinement** |  |  |  |  |
| Rcryst | - | 0.188 | 0.208 | 0.294 |
| Rfree | - | 0.246 | 0.264 | 0.326 |
| **Model composition** |  |  |  |  |
| Non-hydrogen atoms (N) | - | 9544 | 9586 | 6781 |
| Protein residues (N) | - | 1267 | 1267 | 1090 |
| Waters (N) | - | 231 | 241 | 0 |
| Ligands (N) | - | 0 | 2 | 0 |
| **B factors (Å<sup>2</sup>)** |  |  |  |  |
| Protein | - | 45.29 | 55.77 | 148.2 |
| Water/Ligand | - | 61.90 | 84.14 | - |
| **R.m.s. deviations** |  |  |  |  |
| Bond lengths (Å) | - | 0.008 | 0.006 | 0.004 |
| Bond angles (°) | - | 0.929 | 0.768 | 1.397 |
| **Validation** |  |  |  |  |
| Overall | - | 1.56 | 1.58 | 2.45 |
| Clashscore | - | 6.11 | 6.81 | 18.46 |
| **Ramachandran plot** |  |  |  |  |
| Favoured (%) | - | 96.5 | 96.76 | 83.56 |
| Allowed (%) | - | 2.9 | 1.98 | 11.83 |
| Disallowed (%) | - | 0.60 | 0.26 | 4.61 |

In the crystal structure, DOCK10^DHR2^ self-associates through lobe A and forms a complex with CDC42 with 1:1 stoichiometry (**Fig. 3A**). Lobe B contains two antiparallel β sheets and lobe C is formed by a four-helix bundle. Lobe A stabilizes the DOCK10^DHR2^ domain through extensive contacts with lobe B, and as previously mentioned, creates a rigid core. Lobes B and C contact CDC42 to form the catalytic center (**Fig. 3B**), very similar to the DOCK9^DHR2^-CDC42 complex and DOCK2^DHR2^-RAC complexes (Yang et al., 2009; Kilkurni et al., 2011). The major interaction surface of CDC42 with DOCK10^DRH2^ involves its switch 1 and switch 2 regions, which are the conserved nucleotide contact regions of GTPases.

**Figure 3.**
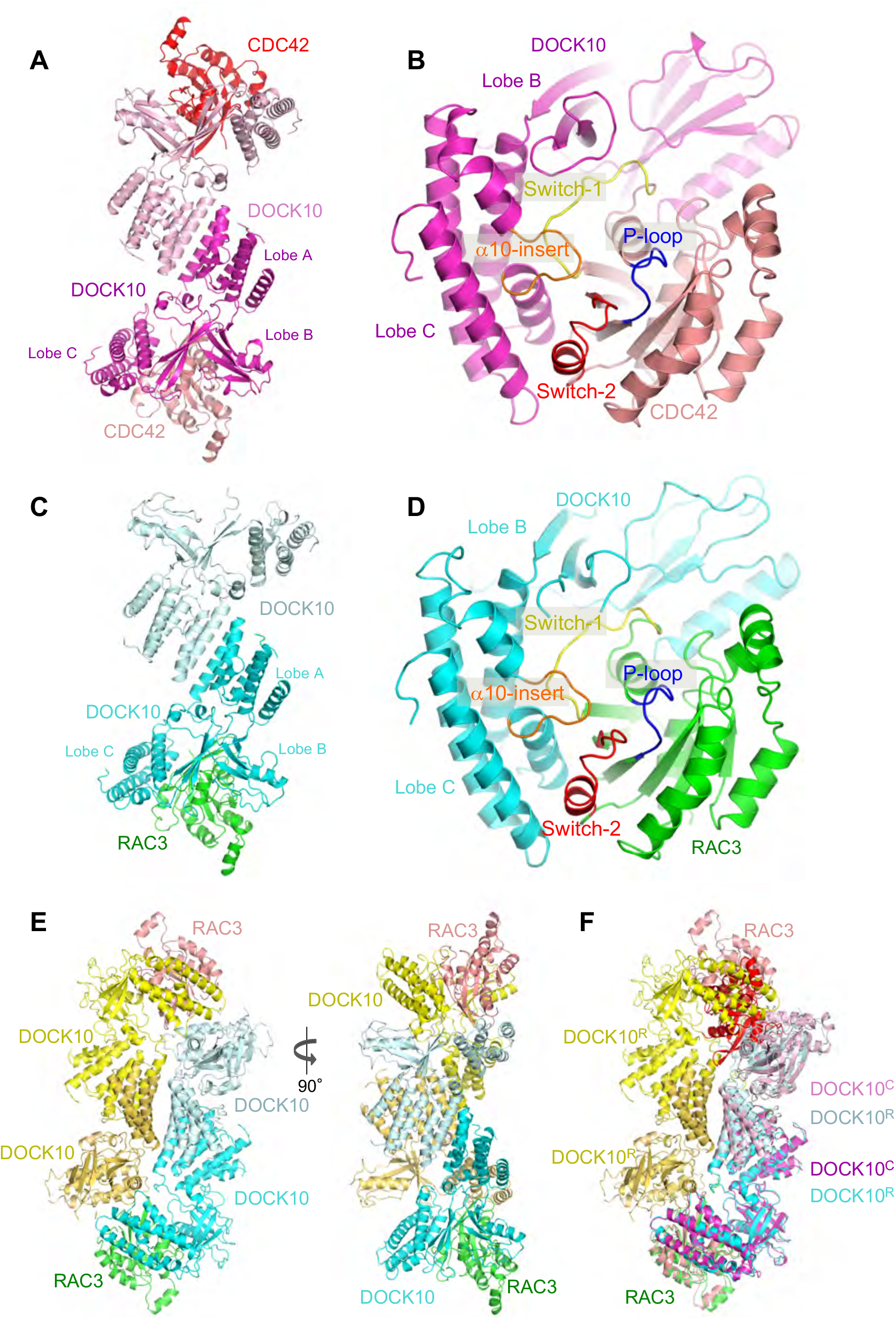
Structures of DOCK10^DHR2^-CDC42 and DOCK10^DHR2^-RAC3. **A**. DOCK10^DHR2^-CDC42 dimer. **B**. Details of CDC42 interactions with the catalytic site of DOCK10^DHR2^. CDC42 interacts with lobs B and C of DOCK10^DHR2^. **C**. DOCK10^DHR2^-RAC3 dimer. **D**. Details of RAC3 interactions with the catalytic site of DOCK10^DHR2^. **E**. Two orthogonal views of the DOCK10^DHR2^-RAC3 tetramer. **F**. Superimposition of the DOCK10^DHR2^-CDC42 dimer onto the DOCK10^DHR2^-RAC3 tetramer.

In contrast to previous DOCK^DHR2^-GTPase structures (Yang et al., 2009; Kilkurni et al., 2011), the DOCK10^DHR2^-CDC42 complex deviates from perfect two-fold symmetry. This is evident by superimposing lobes A and B of mol1 of DOCK10^DHR2^ onto their counterparts in mol2, which indicates asymmetry at the dimer interface (**Supplementary Fig. S2A**). Asymmetry is also observed in the relative orientations of lobe C relative to lobes A and B (**Fig. 4A**). Relative to mol1, in mol2, lobe C, and the associated GTPase, is rotated by a few degrees.

**Figure 4.**
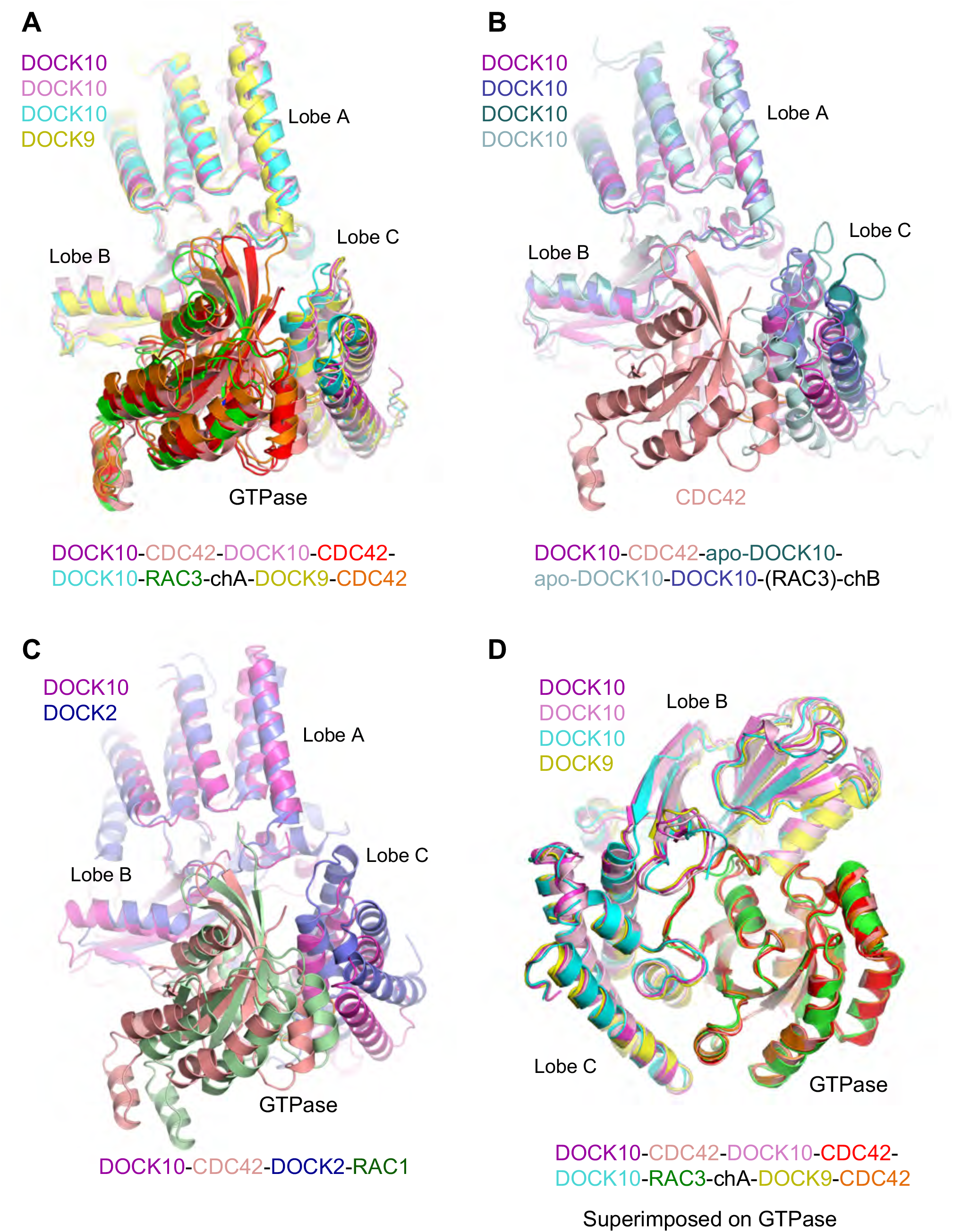
Comparison of DOCK^DHR2^-GTPase complexes. **A**. Superimposition of both DOCK10^DHR2^-CDC42 protomers onto chain A of DOCK10^DHR2^-RAC3 and DOCK9^DHR2^-CDC42 (PDB: 2WM9) (superimposed on lobes A and B). **B**. Superimposition of DOCK10^DHR2^-CDC42 onto both apo-DOCK10^DHR2^ subunits and the GTPase-free DOCK10^DHR2^ domain of the DOCK10^DHR2^-RAC3 complex. **C**. Superimposition of DOCK10^DHR2^-CDC42 onto DOCK2^DHR2^-RAC3 (PDB: 2YIN). **D**. Superimposition of DOCK10^DHR2^-CDC42-DOCK10^DHR2^-CDC42-DOCK10^DHR2^- RAC3-chA-DOCK9^DHR2^-CDC42 onto their cognate GTPases.

By using a crystal-soaking approach, similar to that described for DOCK9^DHR2^-CDC42 and DOCK2^DHR2^-RAC (Yang et al., 2009; Kilkurni et al., 2011), the structure of DOCK10^DHR2^-CDC42 with bound GDP was determined at 2.6 Å (**Supplementary Fig. S3 and Table 1**). This structure shows the same overall fold as the crystal structure of DOCK10^DHR2^-CDC42. The electron density for the CDC42-bound GDP was clearly defined (**Supplementary Fig. S3B**). In contrast to the structure of GDP-bound CDC42, no extra density was found in the ternary structure for Mg^2+^ coordinated to the β-phosphate of GDP, similar to the DOCK9^DHR2^-CDC42-GDP complex (Yang et al., 2009). Thus, these data are in agreement with the magnesium exclusion mechanism previously proposed for DOCK9^DHR2^ (15). Protrusion of the α10 helical insert into the CDC42 guanine nucleotide-binding site allows the invariant Val residue (Val1935 of DOCK10) of the nucleotide sensor (15) to directly occlude the nucleotide-coordinated Mg^2+^. Mg^2+^ enhances nucleotide affinity by neutralizing the negatively charged phosphate groups, so its exclusion would dramatically reduce nucleotide affinity. This correlates to the first step of the exchange reaction, when guanine nucleotide dissociates from CDC42.

### Structural analysis of the DOCK10^DHR2^ domain in complex with RAC3

Stable complexes of DOCK10^DHR2^ with both RAC1 and RAC3 were purified. DOCK10^DHR2^ co-migrates with RAC1 and RAC3 on gel filtration, confirming the stable binding of DOCK10^DHR2^ to RAC1 and RAC3.

The structure of DOCK10^DHR2^ in complex with nucleotide-free RAC3 was determined at 3.5 Å resolution **(Fig. 3C-F and Table 1)**. The crystal structure of the DOCK10^DHR2^-RAC3 complex was immediately intriguing because there is only one molecule of RAC3 bound to the DOCK10^DHR2^ dimer in the asymmetric unit, indicating that DOCK10^DHR2^ and RAC3 form a complex with 2:1 stoichiometry. Whether this unusual stoichiometry is a consequence of crystallization or is a real representation of the solution state was further investigated using SEC-MALS **(Fig. 5A)**. The experimental molecular masses of DOCK10^DHR2^-RAC3 and DOCK10^DHR2^-RAC1 closely match a stoichiometry of 2:1, that is a complex of two subunits of DOCK10^DHR2^ and one molecule of RAC1/3 **(Fig. 5B**), in agreement with the crystal structure.

**Figure 5.**
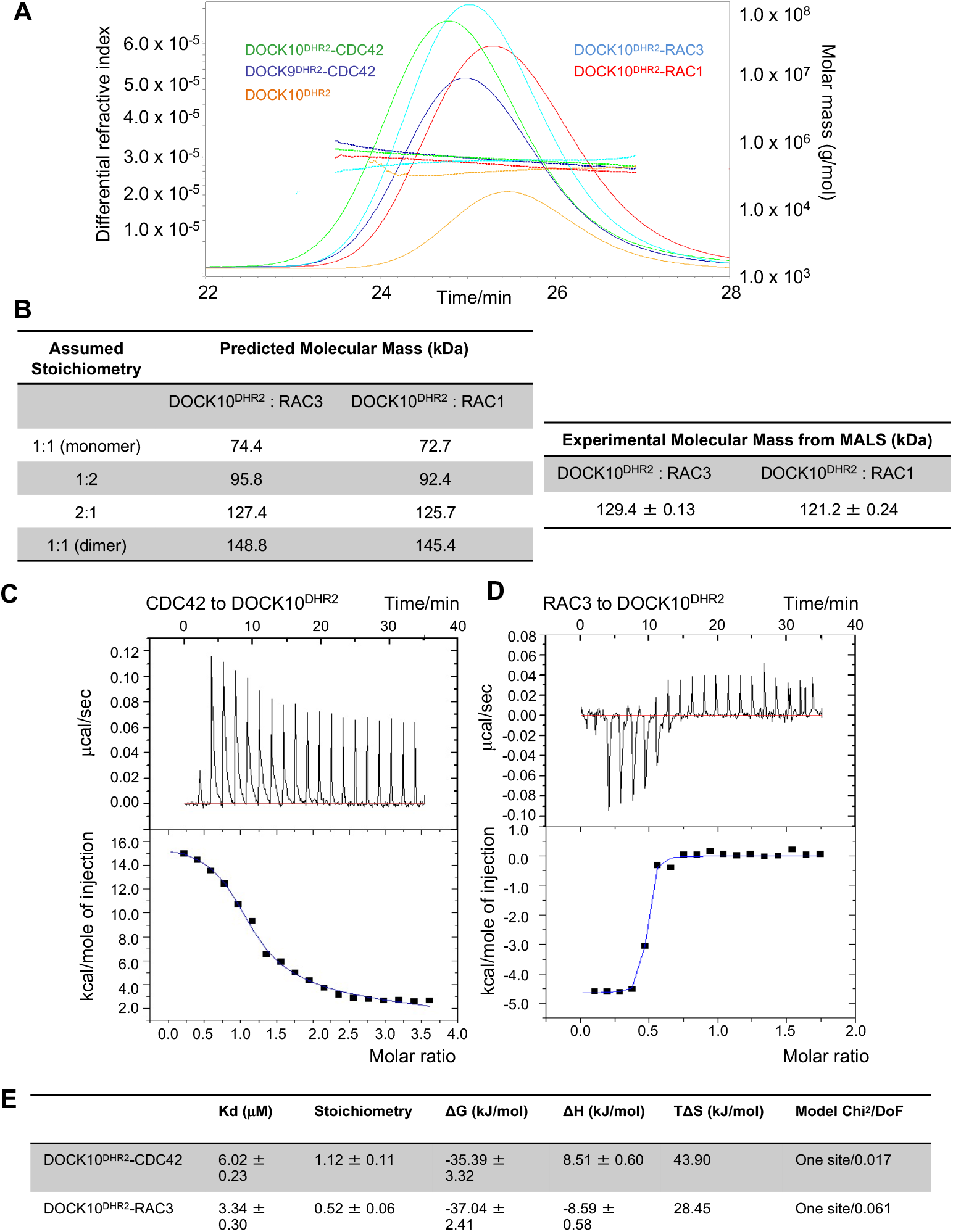
Characterization of interaction between DOCK^DHR2^ proteins with their cognate GTPase. **A**. Size exclusion chromatography/multi-angle light scattering (SEC/MALS) profiles of DOCK^DHR2^ proteins or protein complexes. Molecular masses of DOCK^DHR2^ protein alone and in complex with GTPases were measured with multi-angle light scattering and are plotted against elution time from the 10/300 GL gel-filtration column. Peaks are visualized as unit-less relative refractive index measurements. **B**. Table of molecular mass values obtained from SEC/MALS and the known molar mass for each sample. **C**. Top: Recorded raw ITC data for the DOCK10^DHR2^-CDC42 interaction in black, and the baseline is drawn in red. The bottom panel shows, after the correction by subtraction of HOD, the integrated areas of each injection plotted (black squares) against the molar ratio of titrated CDC42. The blue curve represents the non-linear best fit to the data assuming a one-site binding model. 10 µM DOCK10^DHR2^ was in the cell and 200 µM CDC42 was in the syringe. **D**. Top: Recorded raw ITC data for DOCK10^DHR2^-RAC3 interaction in black, and the baseline is drawn in red. The bottom panel shows, after the correction by subtraction of HOD, the integrated areas of each injection plotted (black squares) against the molar ratio of titrated RAC3. The blue curve represents the non-linear best fit to the data assuming a one-site binding model. 20 µM DOCK10^DHR2^was in the cell and 200 µM RAC3 was in the syringe. **E**. Table of thermodynamic parameters fitted by Origin-ITC for DOCK10^DHR2^ and its GTPase binding. Kd is derived from the binding constant K by the equation K_d_=1/K. Chi-Square = (1/DoF) * S((data value - function value)^2^), where DoF (degree of freedom) = number of points - number of parameters. Chi-Square quantifies the deviation of predicted values from the actual values. If there is no deviation (perfect fit), the Chi-Square equals 0.

In the DOCK10^DHR2^-RAC3 complex, two subunits of DOCK10^DHR2^ self-associate through the canonical interface of lobe A. RAC3 binds to one subunit of DOCK10^DHR2^ similar to CDC42, through contacts with lobes B and C (**Fig. 3A-D**). The major interaction surface of RAC3 involves the switch 1 and switch 2 regions (**Fig. 3D**). The conformation of the DOCK10^DHR2^-RAC3 subunit is essentially similar to that of DOCK10^DHR2^-CDC42 and DOCK9^DHR2^-CDC42 (15) **(Fig. 4A)**. There are small variations in the relative orientations of lobe C, and associated GTPase, with lobes A and B. The structural variation of lobe C is more pronounced when comparing DOCK10^DHR2^-CDC42 and DOCK2^DHR2^-RAC3 (**Fig. 4C**). Thus, DOCK10^DHR2^ catalyzes nucleotide exchange in RAC3 through a similar mechanism to DOCK10^DHR2^ mediated-activation of CDC42.

In the unliganded subunit of DOCK10^DHR2^ in the DOCK10^DHR2^-RAC3 complex, compared with the DOCK10^DHR2^ subunit bound to RAC3, lobe C has rotated away from lobes A and B, creating a less compact structure (**Fig. 4B**). This conformation is similar to one of the apo-DOCK^DHR2^ subunits, where lobe C has rotated away from lobes A and B, making it unable to form stable interactions with a RAC3 molecule. In contrast, in the second DOCK10^DHR2^ subunit of apo-DOCK10^DHR2^, lobe C is positioned closer to lobes A and B, blocking access of the GTPase to its binding sites on lobes B and C (**Fig. 4B**).

Comparing all known structures of the DOCK^DHR2^ domains of the DOCK-D subfamily by superimposing on their cognate GTPase, shows that for the DOCK9^DHR2^-CDC42, DOCK10^DHR2^-CDC42 and DOCK10^DHR2^-RAC3 complexes, the cognate GTPase interacts with lobe C in an essentially identical manner, with a slight shift in the position of lobe B (**Fig. 4D**).

Analysis of the crystal lattice revealed that a dimer of DOCK10^DHR2^-RAC3 dimers associate to form a heterotetramer, whereby a DOCK10^DHR2^-RAC3 protomer interacts with a GTPase-free DOCK10^DHR2^ subunit (**Fig. 3E**). RAC3 of the canonical DOCK10^DHR2^-RAC3 protomer forms additional interactions with lobe C of the GTPase-free DOCK10^DHR2^ subunit through the β2-β3 loop and C-terminal α5 helix. Formation of this heterotetramer is contingent on a GTPase-free DOCK10^DHR2^ subunit, and thus is incompatible with dimers of the DOCK10^DHR2^-CDC42 dimer (**Fig. 3F**).

### Characterization of the interaction between the DOCK10^DHR2^ domain and its cognate GTPases

ITC experiments were performed to further characterize the interactions between DOCK10^DHR2^ and its cognate GTPases. After integrating each peak and subtracting from the heat of dilution (HOD) (**Supplementary Fig S4**), an interaction curve was fitted using the software Origin 7-ITC (**Fig. 5C and D**). Subsequently all the thermodynamic parameters were calculated (**Fig. 5E**) (29,30). The ITC data confirm the stoichiometry of 1:1 between DOCK10^DHR2^ and CDC42 in the DOCK10^DHR2^-CDC42 complex and the stoichiometry of 2:1 between DOCK10^DHR2^ and RAC3 in the DOCK10^DHR2^-RAC3 complex. The binding event between DOCK10^DHR2^ and CDC42 is endothermic. The interaction of DOCK10^DHR2^ and CDC42 is enthalpically unfavorable with a positive ΔH value (ΔH of 8.51 ± 0.06 kJ/mol), but the binding is entropically driven by a large favorable entropy change of (TΔS of +49.3 kJ/mol). Whereas the binding event between DOCK10^DHR2^ and RAC3 is exothermic. The interaction of DOCK10^DHR2^ and RAC3 is enthalpically favorable with a negative ΔH value (ΔH of -8.59± 0.58 kJ/mol), and the binding is also entropically driven by a large favorable entropy change of (TΔS of +28.45 kJ/mol). The enthalpy change of binding reflects the loss of protein–solvent hydrogen bonds and van der Waals interactions, formation of protein–ligand bonds, salt bridges and van der Waals contacts, and solvent reorganization near protein surfaces (31). These individual components produce unfavorable enthalpic contributions to the binding between DOCK10^DHR2^ and CDC42 and favorable enthalpic contributions to the binding between DOCK10^DHR2^ and RAC3.

We suspected that the 2:1 stoichiometry of DOCK10^DHR2^ to RAC3 is caused by a negative cooperative effect mediated by communication from the RAC3-bound DOCK^DHR2^ monomer to the RAC3-free DOCK2^DHR2^ subunit through the DOCK10^DHR2^ dimer interface mediated by lobe A. Based on the structure of the DOCK10 dimer interface we hypothesized that mutating three interfacial residues (highlighted in the **Supplementary Fig. S5A)** would disrupt the hydrophobic network and hence DOCK10 dimerization. The dimerization interface of DOCK10^DHR2^ (lobe A) is remote from the GTPase-binding sites and would not be expected to directly affect GTPase-binding (15). A dimerization-defective triple mutant F1846A/F1853A/F1860A of DOCK10^DHR2^ (DOCK10^DHR2^DM) was generated.

DOCK10^DHR2^DM was incubated with excess RAC3 and analyzed using SEC-MALS in identical conditions as that for the wild type DOCK10^DHR2^. A single symmetric peak in the chromatogram suggests a homogeneous species of DOCK10^DHR2^DM-RAC3. The experimental molecular weight for this species matches the theoretical calculated molecular mass for monomeric DOCK10^DHR2^ bound to RAC3 (**Supplementary Fig. S5B**). There is no extra peak in the chromatography trace accounting for apo-DOCK10^DHR2^DM, although a peak representing excess RAC3 was observed. This suggested that all monomeric DOCK10^DHR2^DM bound RAC3, and supports the idea that RAC3 binds to dimeric DOCK2^DHR2^ through negative cooperativity.

### Structural basis for the specificity of DOCK10 for both CDC42 and RAC3

CDC42 and RAC3 contact DOCK9 and DOCK10 through two segments: residues 1-74 (segment 1, incorporating switch 1, the P-loop and switch 2) and 159-170 (segment 2). Within segment 1, RAC3 and CDC42 differ by 13 residues, eight of which contact DOCK9 and DOCK10 (T3A (β1), K27A (α1), S30G, V33I (switch 1), A41S (β2), T43N (β2), T52N and F56W (switch 2, β3)) (CDC42 amino acid – sequence number – RAC3 amino acid). In segment 2, all contact residues are conserved between CDC42 and RAC3. Thus, to explain the differences in specificities between DOCK9 and DOCK10 for CDC42 and RAC3, we focus on the variable residues of segment 1 of the GTPases. Six residue differences stand out as likely important in defining the ability of DOCK10 to activate RAC3 because residues of DOCK10 that directly and indirectly contact these residues differ from their counterparts in DOCK9, whereas K27A and S30G contact sites are identical between DOCK9 and DOCK10. The most significant contact differences involve V33I, A41S, T43N and F56W.

Starting with the T43N difference. In RAC3, Asn43 forms a hydrogen bond to Arg1876 of DOCK10 (**Fig. 6A**). This is not possible for DOCK9 where Thr substitutes for Arg1876. In the DOCK10^DHR2^-CDC42 complex, Thr43 of CDC42 forms a water-mediated hydrogen bond to Arg1876 of DOCK10 (**Fig. 6B**). Thus, DOCK10 forms additional hydrogen bonds to CDC42 and RAC3, which are not compensated in DOCK9.

**Figure 6.**
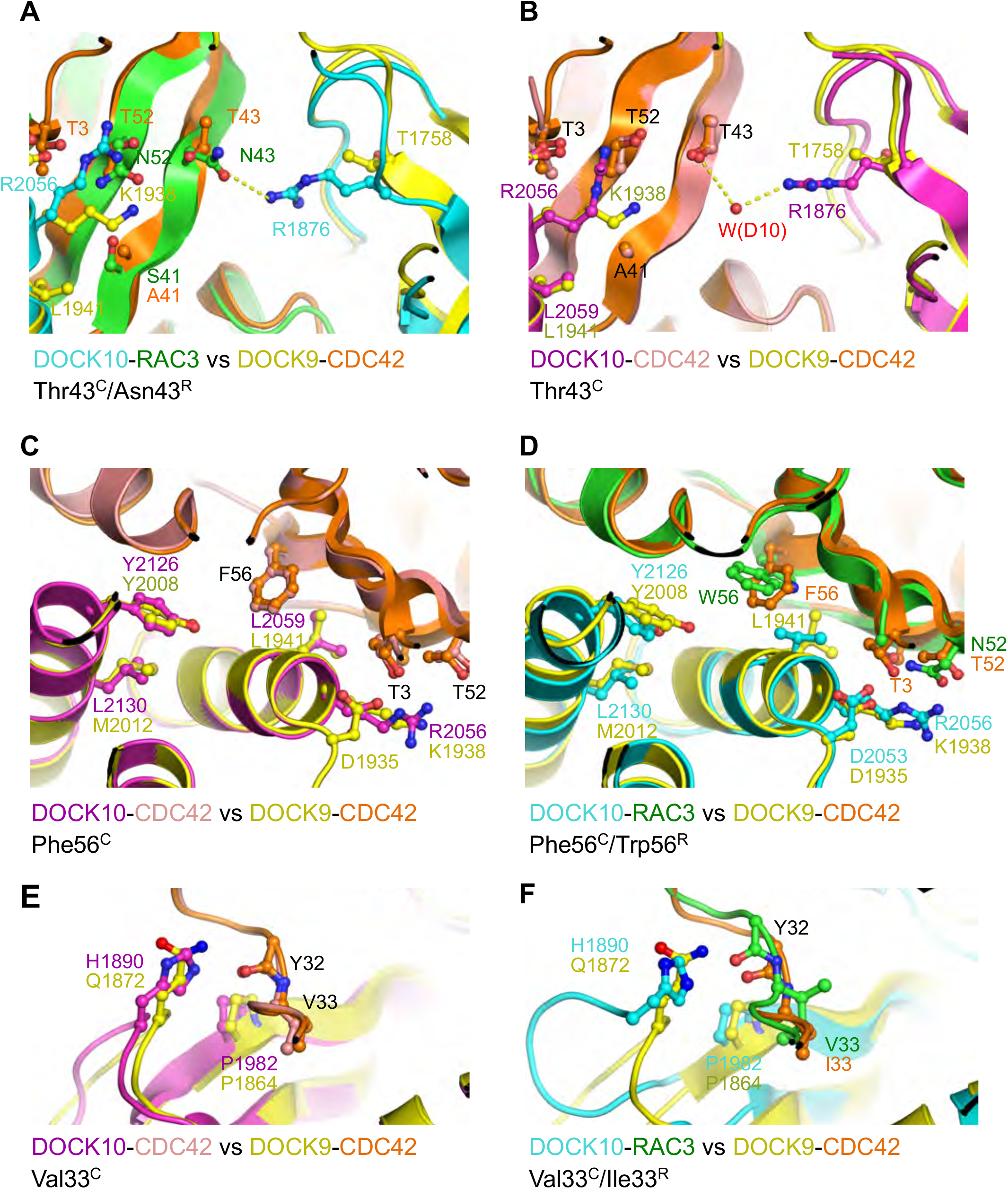
Basis of dual activity of DOCK10^DHR2^ to CDC42 and RAC3. **A and B**. Differences in Thr43 (CDC42) and Asn43 (RAC3) between DOCK10 ^DHR2^-RAC3 compared with DOCK9 ^DHR2^-CDC42 (**A**) and DOCK10 ^DHR2^-CDC42 compared with DOCK9 ^DHR2^-CDC42 (**B**). **C** and **D**. Differences in Phe56 (CDC42) and Trp56 (RAC3) between DOCK10 ^DHR2^-CDC42 compared with DOCK9 ^DHR2^-CDC42 (**C**) and DOCK10 ^DHR2^-RAC3 compared with DOCK9 ^DHR2^-CDC42 (**D**). **E** and **F**. Differences in Val33 (CDC42) and Ile33 (RAC3) between DOCK10 ^DHR2^-CDC42 compared with DOCK9 ^DHR2^-CDC42 (**E**) and DOCK10 ^DHR2^-RAC3 compared with DOCK9 ^DHR2^-CDC42 (**F**).

A second major difference is the substitution of the bulky Trp56 in RAC3 for Phe56 in CDC42. In both the DOCK9-CDC42 and DOCK10-CDC42 complexes, the aromatic side chain of Phe56 of CDC42 is surrounded by the same residues, notably Leu1941 (2059 in DOCK10) and Tyr2008 (2126 in DOCK10) (**Fig. 6C**). The conformation of Phe56^CDC42^ and its neighboring residues is identical in both DOCK9-CDC42 and DOCK10-CDC42 complexes. To accommodate the bulkier Trp56 residue of RAC3, its side chain rotates, causing concomitant movement of Leu2059^DOCK10^ and Tyr2126^DOCK10^ relative to their positions in the DOCK9-CDC42 complex (**Fig. 6D**). The position of the Leu1941/2059 side chain in the DOCK9/10-CDC42 complexes, which is not compatible with a Trp at position 56, is prevented in RAC3 due to a close contact with Ser41 (Ala in CDC42) (**Fig. 7B**). Thus, the position of Leu2059^DOCK10^ required for RAC3 interactions requires a Ser at position 41. Although this does not provide specificity directly, it is of note that Ser41 participates in a network of hydrogen bond involving Asn52 and Asn43, the latter forming a hydrogen bond to Arg1876^DOCK10^ (described above) (**Fig. 7**). Arg2056^DOCK10^ (Lys1938 in DOCK9) therefore may play a role in stabilizing the conformation of this hydrogen bond network, thus allowing DOCK10 and not DOCK9 to bind RAC3.

**Figure 7.**
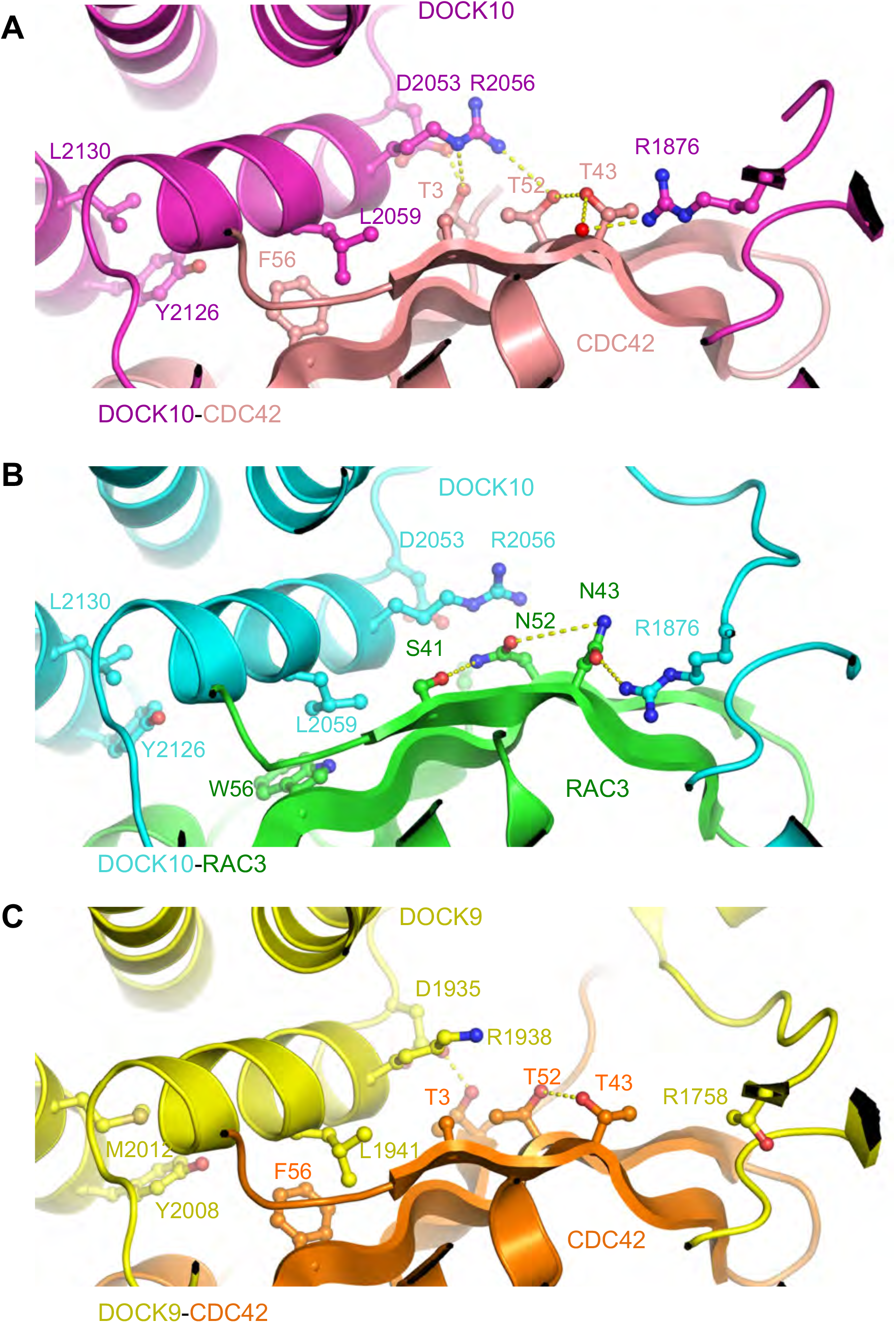
Basis of dual activity of DOCK10^DHR2^ to CDC42 and RAC3. **A**. Network of H-bonds involving S41^RAC3^ contacting R1876^DOCK10^ in the DOCK10^DHR2^-RAC3 complex, absent from DOCK10^DHR2^-CDC42 and DOCK9^DHR2^-CDC42. DOCK10 ^DHR2^-CDC42. **B**. DOCK10^DHR2^-RAC3. **C**. DOCK9 ^DHR2^-CDC42.

Another structural change between DOCK9 and DOCK10 possibly important in accommodating the bulky Trp56 residue is that in the DOCK10-RAC3 complex, Tyr2126^DOCK10^ shifts to make way for the bulky Trp56^RAC3^ side chain (**Fig. 6C, D**). This is possible in DOCK10 because Trp2126^DOCK10^ packs against Leu2130^DOCK10^. The equivalent Met 2012 in DOCK9 would potentially prevent movement of Tyr2008^DOCK9^.

Finally, in RAC3, the buried Ile33^RAC3^ side chain instead of Val33 of CDC42 pushes the switch 1 region away from the central β-sheet closer to DOCK10. The main-chain carbonyl of Tyr32 whose position is altered contacts His1989 in DOCK10 and Gln1872 in DOCK9 (**Fig. 6E, F**). Possibly, the geometry of the hydrogen bond between His1989 to the main-chain carbonyl of Tyr32 in the DOCK10-RAC3 complex is more favorable than that provided by Gln1872 of DOCK9.

## Discussion

It was generally accepted that the DOCK family GEFs have different specificities towards GTPases: DOCK-A and DOCK-B are RAC specific, DOCK-D are CDC42 specific, and DOCK-C are dual-specific for CDC42 and RAC (1,2). The specificity of different subfamilies of DOCK proteins was assigned based on extrapolation of assays performed on a single intensively studied subfamily member. For example, DOCK-D family GEFs were assumed to be CDC42-specific because DOCK9^DHR2^ was shown to specifically activate CDC42 both *in vitro* and in *vivo*, but not RAC1 or RHOA. However, our pull-down analysis and activity assays showed an unexpected binding and GEF activity of DOCK10 towards RAC (**Fig. 2**). While preparing this study for publication two studies reported findings that DOCK10 activates both CDC42 and RAC (10,11). Furthermore, we found that DOCK7^DHR2^ of the DOCK-C subfamily was CDC42 specific, contrary to the dual specificity proposed from *in vivo* studies (18,19). (20) reported recently *in vitro* a two-fold stimulation of nucleotide exchange of RAC1 by a sub-domain of DOCK7^DHR2^ comprising only the B and C lobes.

In order to understand how this specificity towards different GTPases is achieved at the molecular level, structural studies of the dual-specific DOCK10^DHR2^ were performed. The crystal structure of DOCK10^DHR2^-RAC3 was determined, combined with SEC-MALS analysis and ITC data, confirmed that in solution DOCK10^DHR2^ forms a complex with RAC1/3 with a unique stoichiometry of 2:1 (**Fig. 5**). This leads to another interesting question, namely how binding of one molecule of RAC3 to DOCK10^DHR2^ prevents binding of a second RAC3 to the other DOCK10^DHR2^ subunit of the dimer. Some clues towards answering this question were obtained when monomeric DOCK10^DHR2^ was shown to bind RAC3 with equal stoichiometry (**Supplementary Fig. S5B**). This suggests that dimerization of DOCK10^DHR2^ allows for negative cooperativity between the two catalytic sites of the dimer, mediated by RAC1/3-induced asymmetry of the DOCK10^DHR2^ dimer.

The specificity of GEF towards RHO GTPases is determined by their ability to directly interact with these GTPases. Despite their high sequence identity, DOCK10^DHR2^ displays activity towards RAC, whereas DOCK9 and DOCK11 do not (this work and (11)). To explore the factors underlying the selectivity of DOCK9 and DOCK11 towards CDC42 versus RAC, we began by evaluating the residues that contact the GTPases. Non-conserved residues forming the DHR2 domain interface with the GTPase are likely candidates for determining specific selectivity.

While detecting protein interactions *in vitro* is important for understanding protein function, it is equally important to be able to analyze protein interactions *in vivo*. A prior study had noted that levels of GTP-bound RAC1 increased in the DOCK10 depleted cell lines (5). The reason for the discrepancy between our finding and the elevated active RAC1 remains unclear. First of all, this study has focused on the DOCK10^DHR2^ domain. The N-terminal of DOCK10^DHR2^ may regulate the GEF activity of DOCK10^DHR2^ towards RAC3 and CDC42 differently *in vivo*. Secondly, RAC1 can be activated by many different GEFs and there is functional redundancy between closely related DOCK GEFs. We cannot exclude the possibility that knockdown of DOCK10 indirectly enhances the capability of other GEFs to activate RAC.

Our biochemical data supported by structural analysis report a detailed characterization of a putative interaction between DOCK10^DHR2^ and RAC1/3. DOCK10^DHR2^ domain can directly interact with nucleotide-free RAC and CDC42 *in vitro* and catalyzes the nucleotide exchange. It gives us a full picture of DOCK10 catalytic activity, and would help to reveal DOCK10 mediated signaling pathways in regulating cellular mobility. Additional findings indicate that the dual specific GEF activity towards CDC42 and RAC GTPases is unique to DOCK10 within its evolutionarily related subfamily. The marked differences between the specificity toward GTPases amongst DOCK9, DOCK10 and DOCK11 are intriguing because the DHR2 domain of these three DOCK proteins share ∼66 % amino-acid identity (**Supplementary Fig. S1**).

Furthermore, DOCK10^DHR2^ is now understood in molecular details from multiple structural perspectives. However, *in vitro* conditions do not fully recapitulate the physiological conditions. The goal for the future will be to understand how the GEF activity of DOCK family members is integrated into the complex signaling networks that control fundamentally important processes in cell biology. In addition, our *in vitro* functional and structural studies provide novel information when designing specific inhibitors of a particular DOCK GEF, which could assist the targeted interruption of a DOCK GEF activity *in vivo* to better understand cellular signaling pathways.

## Experimental Procedures

### Cloning and mutagenesis

Guided by the GEF catalytic activity and CDC42 binding data of their family members, the DHR2 domain boundaries of DOCK10, DOCK11 and DOCK7 were defined by sequence conservation (**Supplementary Fig. S1**) (15,17-19). Human DOCK10 DNA was synthesized by Clontech (NM_014689). The DHR2 domain of human DOCK11 DNA (NM_144658.3) was synthesized by BioMatik (USA). The human DOCK7 DNA (NM_033407.2) was a gift from Professor Linda van Aelst at Cold Spring Harbor Laboratory. To screen the DHR2 domain for expression, solubility and suitability for crystallization, several constructs were generated that differ at their N-termini. All constructs were cloned into the pOPNIN-SUMO* vector encoding a N-terminal His_6_-SUMO tag and a C-terminal double-Strep (DS) tag as described in (15). The soluble and enzymatic active constructs described here include DOCK10^DHR2^ (residues 1695-2151), DOCK7^DHR2^ (residues 1673-2112), and DOCK11^DHR2^ (residues 1609-2073).

RAC3 (residues 1-192) and RHOA (residues 3-180) were amplified by PCR from cDNA library of human heart muscle, and were fused to DNA sequences encoding a N-terminal GST tag. CDC42 (residues 1-191) and RAC1 (residues 1-192) were cloned into pGEX6P. The USER™ (uracil-specific excision reagent) Friendly Cloning Kit (NEB) was used for all the cloning procedures. All sequences were confirmed by GATC Biotech.

### Protein expression and purification

The different DOCK^DHR2^ constructs were freshly transformed into expression strain *E. coli* B834 (DE3) pRare cells and expressed using 0.2 mM IPTG for 16 h at 16 °C. Cells were pelleted, resuspended in buffer A (50 mM Tris.HCl (pH 8.0), 500 mM NaCl, 10% glycerol, and complete EDTA-free protease inhibitor tablets (Roche)), lysed by sonication on ice, and clarified by centrifugation at 20,000 *g* for 30 min at 4 °C. The soluble fraction was purified using either 10 mL nickel-chelating column or Strep-Tactin column (Qiagen). The N-terminal His-SUMO tags were cleaved from the eluted protein by TEV protease overnight and removed by passing through a second nickel column. DOCK^DHR2^ domains were further purified by a HiLoad 26/60 Superdex 75 gel filtration column (GE Healthcare) and a RESOURCE™ Q column (GE Healthcare). GST-tagged GTPases were purified on high affinity GST resin (GenScript). The GST-tags were removed by PreScission protease and separated from the GTPase by a further gel filtration column. 5 mM EDTA was included in the buffer to remove nucleotide. DOCK^DHR2^-GTPase complexes was prepared by incubating the purified DOCK^DHR2^ and GTPase at a 1:2 molar ratio on ice for 1 h, and then separating the complex from free GTPase by a HiLoad 26/60 Superdex 200 gel filtration column (GE Healthcare), followed by a RESOURCE™ Q column. The purity of the proteins was visualized by standard SDS-PAGE techniques.

### Crystallization and structure determination

Crystals of apo-DOCK10^DHR2^ (4.5 mg/mL) were grown in seeded buffer containing 15% (w/v) PEG 20 K, 9% glycerol, 10 mM spermine tetrahydrochloride, 0.1 M Tris-HCl (pH 8.5) at 12 °C by micro-batch crystallization under paraffin oil. Crystals of DOCK10^DHR2^-CDC42 complex (5 mg/mL) were grown in buffer containing 19% (w/v) PEG 2000 MME, 10 mM strontium chloride hexahydrate, 100 mM HEPES (pH 7.5) at 20 °C by the sitting drop vapor diffusion method, in which 0.3 μL of the protein is added to 0.3 μL crystallization buffer in the sample well above 50 μL reservoir solution. For the DOCK10^DHR2^-CDC42-GDP complex, the pre-existing DOCK10^DHR2^-CDC42 crystals were incubated in crystallization buffer supplemented with GDP-MgCl_2_ to a final concentration of 5 mM for 5 min prior to cryo-protection. The native crystals of DOCK10^DHR2^-R1 (6.2 mg/mL) were grown using sitting-drop vapor diffusion methods, in buffer containing 25% (w/v) PEG 3350, 200 mM potassium acetate, 8% (v/v) 1,1,1,3,3,3-Hexafluoro-2-propanol, at 20 °C. Crystals of DOCK10^DHR2^-RAC3 complex (8.1 mg/mL) were grown using sitting-drop vapor diffusion methods, in buffer containing 25% (w/v) PEG 1500, 100 mM MMT, at 20 °C. MMT buffer is produced by mixing DL-malic acid, MES and Tris base in the molar ratios of 1:2:2.

The crystals were either cryo-protected by the precipitant present in their crystallization buffer or incubated in mother liquor with addition of 25% (v/v) glycerol for ∼ 90 s prior to freezing in a nitrogen gas stream at 100 K. The diffraction data were collected at either Diamond Light Source (DLS), processed using iMosflm and scaled using SCALA SCALA (21,22). The initial phase information was obtained by molecular replacement (MR) using PHASER (23). DOCK9^DHR2^-CDC42 structure (PDB 2WM9) and RAC3 coordinates from the RAC3-GDP (PDB 2C2H) were used accordingly as search models. The structure was refined and completed using cycles of PHENIX or REFMAC (22,24,25). Non-crystallographic symmetry restraints and parameters generated by TLSMD sever were applied throughout the refinement. Water molecules were added towards the end of refinement. The structure of apo-DOCK10^DHR2^ was processed at a lower resolution of 5 Å, which meant that the data-to parameter ratio was very low. Therefore additional prior knowledge was applied to the initial model, such as the added deformable elastic network (DEN) restraints from known homologous structures to allow global and local deformations of these homology models (26). MOLPROPBITY was used to check for steric clashes and residue stereochemistry (27). Data collection and refinement statistics are given in **Table 1**.

### Pull-down assays

*In vitro* pull-down experiments were performed with separately purified DOCK DHR2 domain and GTPases to test for direct binding. Equally amounts of Strep-Tactin super flow plus beads (15 μL, Qiagen) were saturated with DOCK^DHR2^ proteins (40 μg). The GTPase was added at a two-fold excess molar ratio. After incubation for 1 h at 4 °C, the beads were washed five times with buffer, boiled and visualized by Coomassie blue staining on a 15% SDS-PAGE gel. Non-specific binding of the GTPases to the beads was also assessed in absence of the Strep-Tagged DOCK proteins.

### Real time kinetics GEF activity assay

All fluorescence intensity measurements were either carried out on a Cary Eclipse (VARIAN) fluorescence spectrophotometer or on a POLARstar Omega microplate reader (BMG Tech). Firstly, 2 μM nucleotide-free DOCK10^DHR2^- GTPase complex was added into a well containing 50 mM Tris.HCl (pH 8.0), 200 mM NaCl, 10 mM MgCl_2_, 2 mM DTT and 2 μM mant-GDP at 20 °C, to initiate mant-GDP binding to the GTPase. The second step is to initiate mant-GDP/GTP exchange on the GTPase with 200 μM GTP. The fluorescence signals (λem=355 nm; λex=460 nm) were monitored every 15 s for 250 cycles. Data were analyzed using GraphPad Prism 5 (*www*.*graphpad*.*com*).

### Small angle X-ray scattering (SAXS)

The experimental SAXS data for DOCK10^DHR2^ were collected in a range of different concentrations (1-8 mg/mL) by the flow mode on the EMBL BM29 beam at 10 °C. 10 frames of the scattering intensity were recorded at 2 s/frame, as a function of the scattering angle (2θ). A similar measurement was performed on the SAXS buffer both before and after the sample for the data correction of DOCK10^DHR2^ scattering. The theoretical scattering from the atomic model of DOCK10^DHR2^ was calculated using CRYSOL and evaluated against the measured scattering data (28).

### Multi angle light scattering (MALS)

Purified protein or protein complex (>1.5 mg/mL) was subject to a 24 mL Superdex 200 analytical gel filtrating column 10/300 GL (GE healthcare) at a rate of 0.5 mL/min at 20 °C by HPLC (Varian). MALS was performed on the elute at 658 nm using 18-angle miniDAWN™ TREOS MALS light scattering detector (Wyatt Technology) attached to a refractive index detector (Wyatt Optilab rEX) and a UV absorbance detector. Signals from the light scattering photometer and the refractometer were processed using the ASTRA software (Wyatt Technology). A plot of K*c/R(θ) vs. sin2(θ/2) was constructed to solve the molecular weight.

### Isothermal titration calorimetry (ITC)

All ITC experiments were performed using a VP-ITC200 microcalorimeter (MicroCal). The purified protein samples were exhaustively dialyzed against the ITC buffer at 4 °C, and then centrifuged at 40 K rpm for 10 min to remove aggregation and degas just before use. The ITC buffer comprised 20 mM Tris.HCl (pH 8.0), 150 mM NaCl, 2 mM DTT, 10% glycerol. The protein sample in the cell was typically at a concentration of approximately 10-20 μM, whereas the sample in the syringe was about 100-200 μM. Experiments were performed with 6 μcal/sec reference power at 500 rpm stirring rate. Each run consists of an initial 0.4 μL injection, followed by 18 injections of 2 μL with 105 or 120 sec intervals between each injection. Data were analyzed using Origin7-ITC, and the baselines were adjusted manually before integrating the area under each peak (29). After subtracting the heat of dilution injections, the resulting data were fitted to a one site-binding model, providing best-fit values of change in enthalpy (ΔH), and the binding constant (Kd). The fits gave a stoichiometry ratio for the complex.

## Acknowledgments

This work was funded by a Cancer Research UK (C576/A14109) grant to D.B. We thank Professor Linda van Aelst (CSHL) for the DOCK7 cDNA.

## Author contributions

DB and DF designed experiments. DF and NC performed crystallography and SAXS, DF and JY performed GEF assays, DB and DF wrote the paper.

---

**Supplementary Figure S1.**
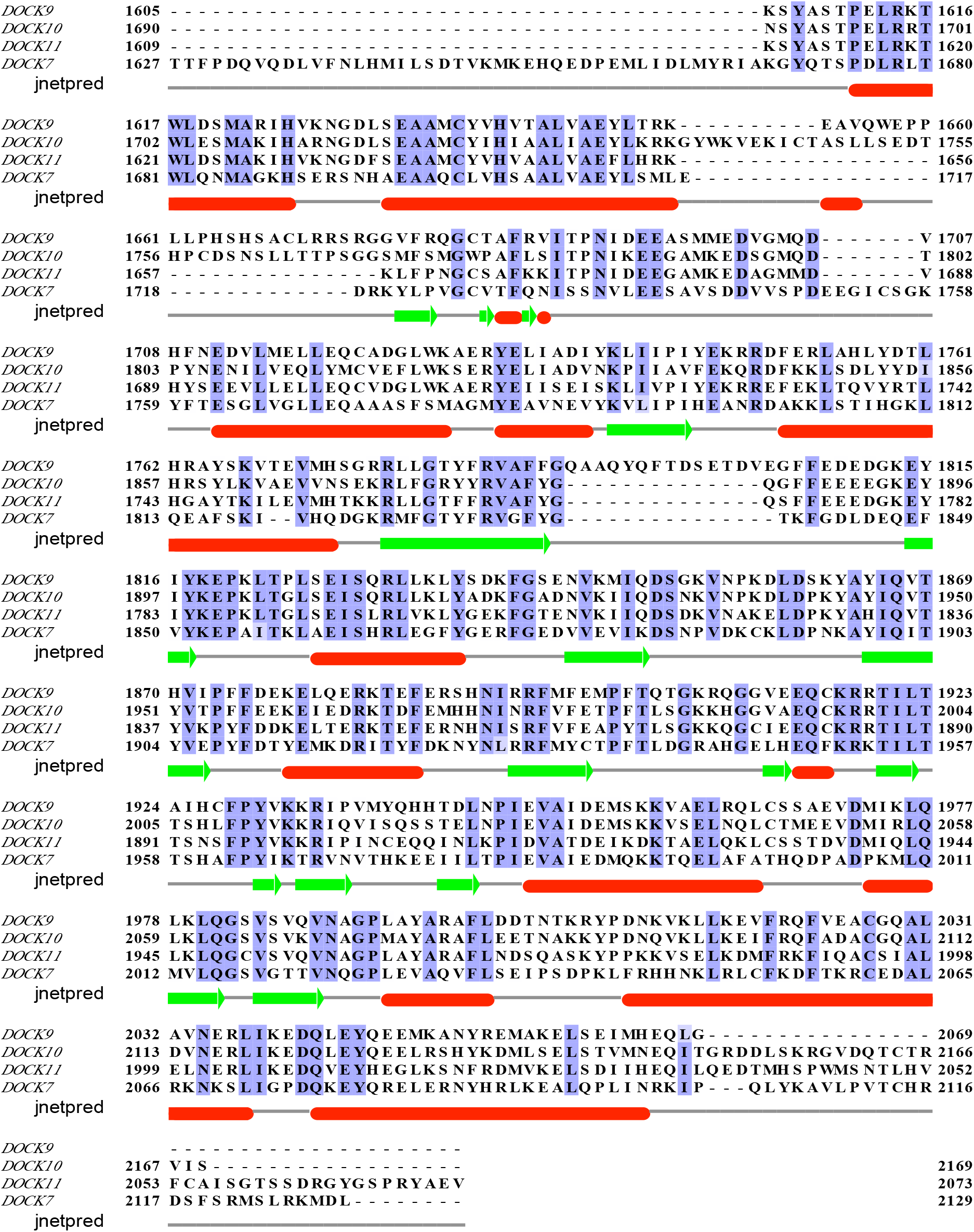
Multiple sequence alignment of the DHR2 domains of DOCK9, DOCK10, DOCK11 and DOCK7. The sequences were aligned using JalView program.

**Supplementary Figure S2.**
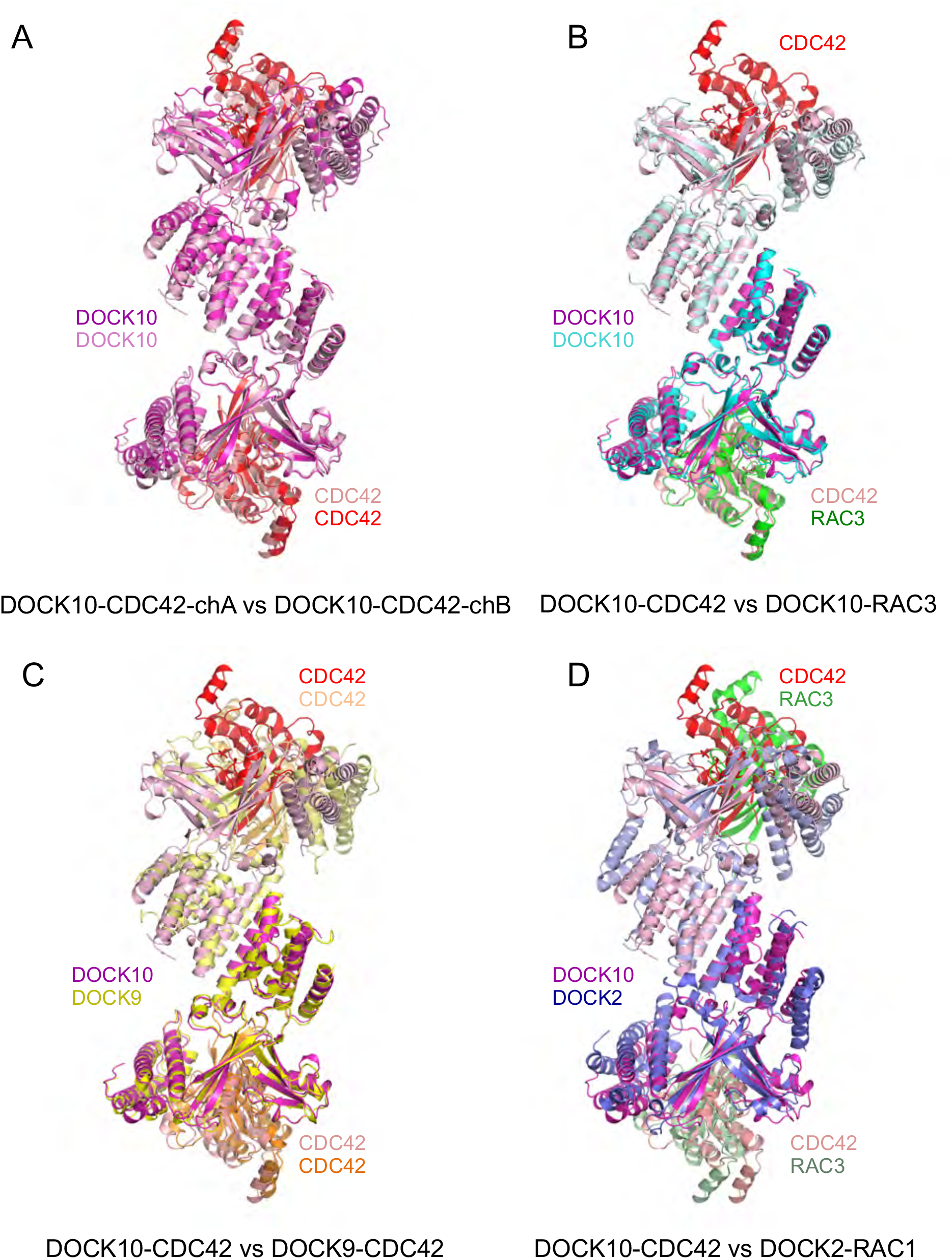
Comparison of DOCK^DHR2^-GTPase dimers. **A**. DOCK10^DHR2^-CDC42-chA vs DOCK10^DHR2^-CDC42-chB. **B**. DOCK10^DHR2^-CDC42 vs DOCK10 ^DHR2^-RAC3. **C**. DOCK10^DHR2^-CDC42 vs DOCK9^DHR2^-CDC42. **D**. DOCK10^DHR2^-CDC42 vs DOCK2^DHR2^-RAC1. All superimposed on lobes A and B.

**Supplementary Figure S3.**
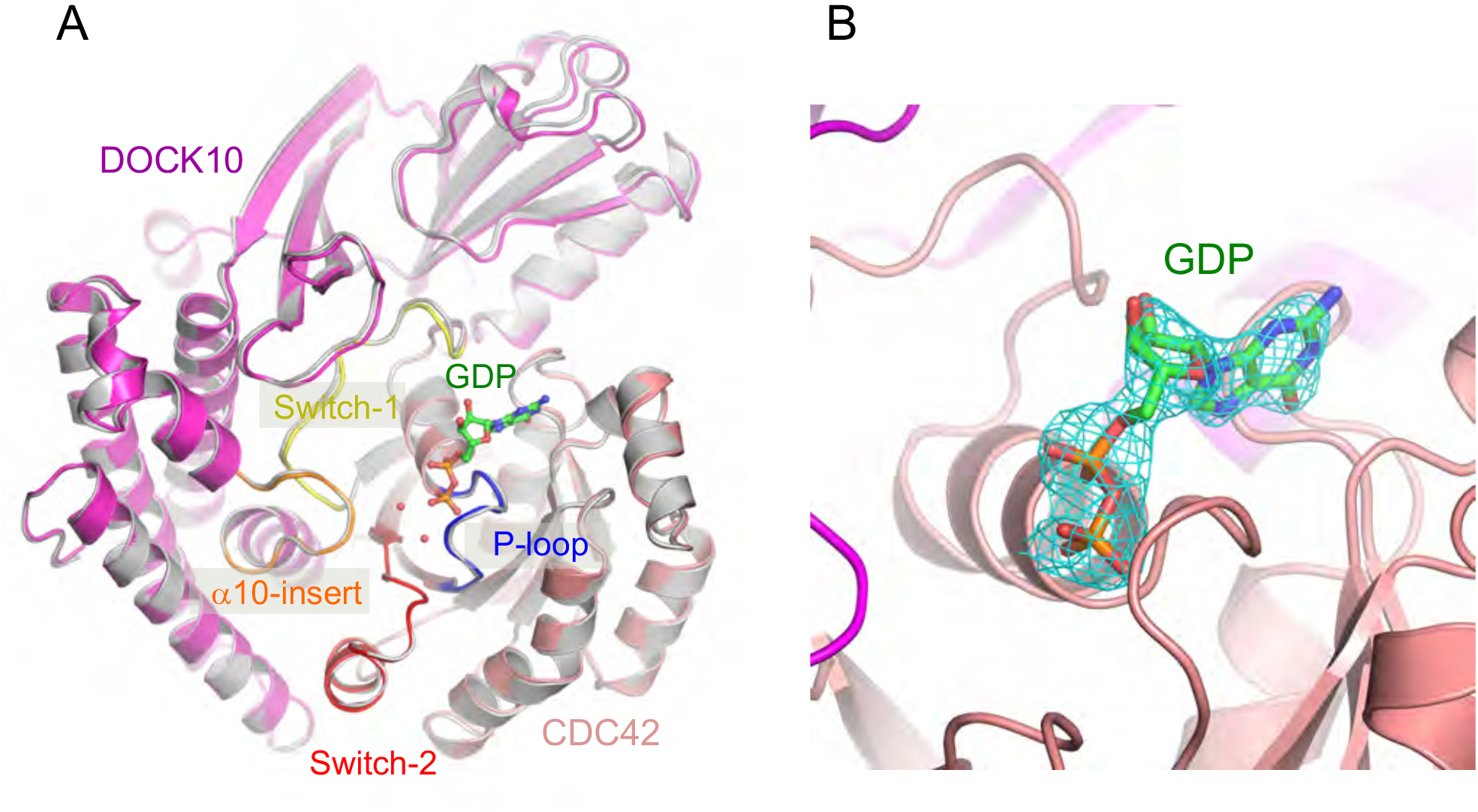
Structure of the DOCK10^DHR2^-CDC42-GDP complex. **A**. Superimposition of the GDP bound *Hs*-DOCK10^DHR2^-CDC42 complex with the nucleotide-free DOCK10^DHR2^- CDC42 complex. In the structure of GDP bound DOCK10^DHR2^-CDC42 complex, DOCK10^DHR2^ is colored cyan and CDC42 is colored wheat. GDP molecule is within the catalytic pocket. Nucleotide-free DOCK10^DHR2^-CDC42 complex is colored in grey. RMSD of 1.72Å between equivalent Cα-atoms of two molecules of DOCK10^DHR2^-CDC42 in presence or in absence of GDP. Global RMSD was calculated by SuperPose Version 1.0. **B**. Details of catalytic center of the DOCK10^DHR2^-CDC42- GDP ternary structure, with atoms of GDP highlighted (red for oxygen atoms, blue for nitrogen atoms and yellow for carbon atoms). Electron density map around GDP is contoured at 2 s (blue) and 4s (green) from a 2F_0_-F_C_ map. Clear density is present for the phosphate.

**Supplementary Figure S4.**
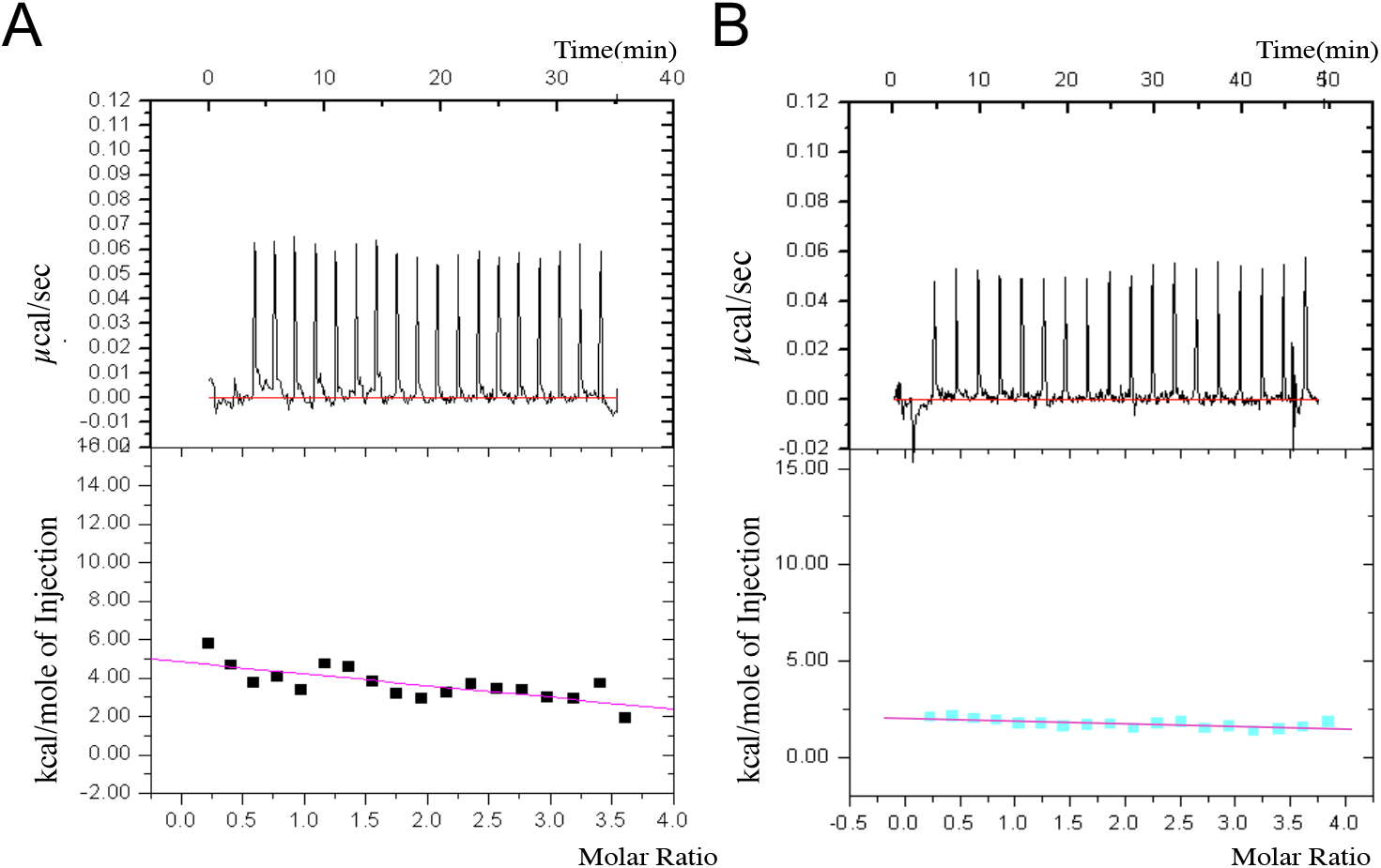
ITC data for the heat of dilution (HOD) of the GTPases of DOCK10. **A** and **B** The top two diagrams show the recorded raw ITC data for heat of dilution (HOD) of CDC42 and RAC3 respectively in black, and the baseline is drawn in red. The bottom two diagrams show the integrated areas of each injection plotted (in black and in cyan against the molar ratio of CDC42 and RAC3 titrated respectively. A linear progression line in purple is fitted for each case.

**Supplementary Figure S5.**
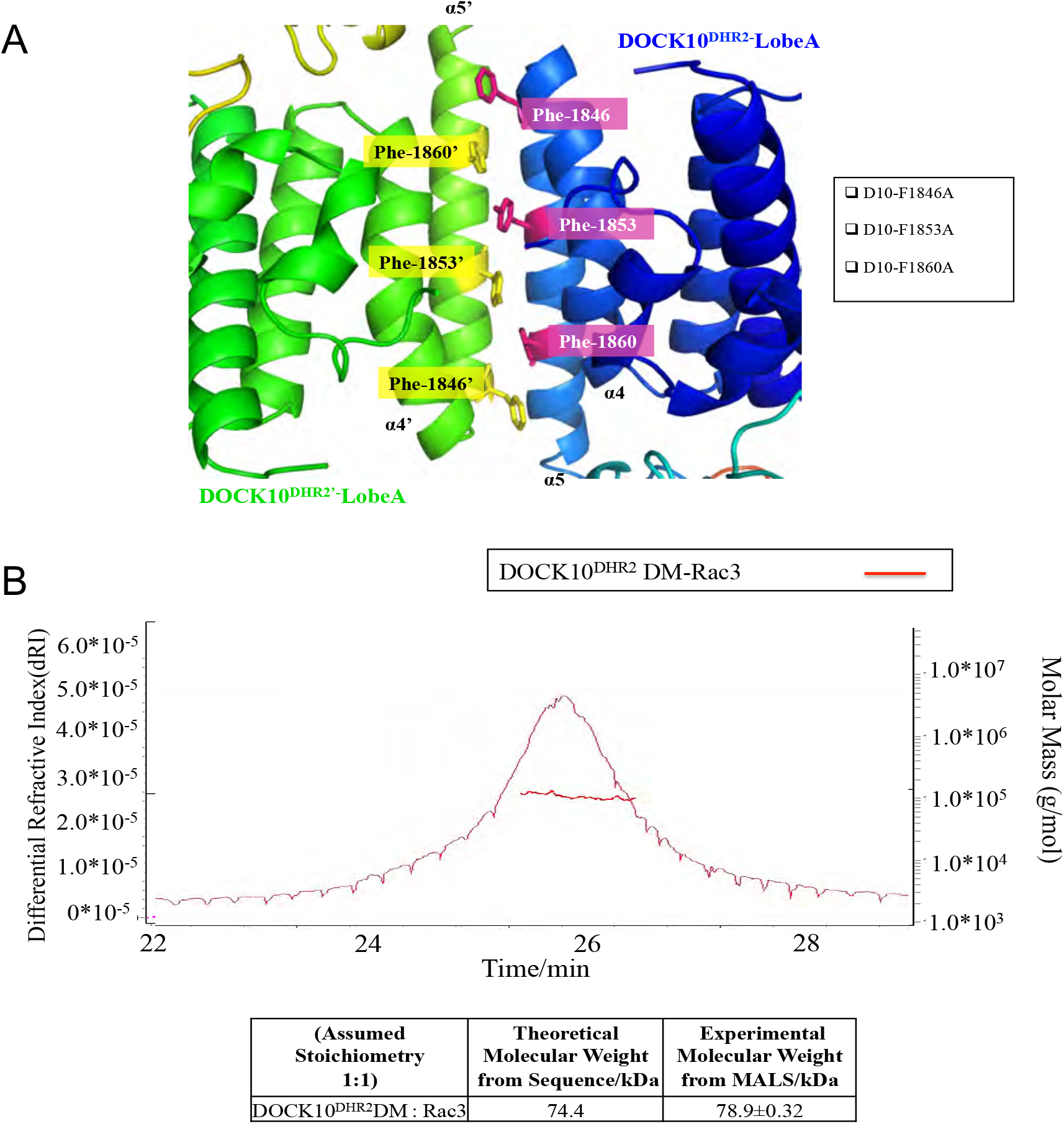
A. Details of the dimer interface. The dimer of DOCK10^DHR2^ formed through the α4 and α5 helices of lobe A. The dimer mutant of DOCK10^DHR2^ is generated by mutating the three aromatic residues highlighted and is designated as DOCK10^DHR2^DM. **B**. A size exclusion chromatography/multi-angle light scattering (SEC/MALS) profile of DOCK10^DHR2^DM-RAC3. (Top) A curve shows refractive index of the protein eluted from the gel filtration column and the dotted line crossing the peak indicates its corresponding molecular mass. Mass estimates are determined for sample over whole elution peak. Bottom: table shown the theoretical and the experimental molecular mass for the monomeric DOCK10^DHR2^DM in complex with RAC3.

## References

1. Cote, J. F., and Vuori, K. (2002) Identification of an evolutionarily conserved superfamily of DOCK180-related proteins with guanine nucleotide exchange activity. Journal of cell science 115, 4901–4913

2. Meller, N., Merlot, S., and Guda, C. (2005) CZH proteins: a new family of RHO-GEFs. Journal of cell science 118, 4937–4946

3. Meller, N., Irani-Tehrani, M., Kiosses, W. B., Del Pozo, M. A., and Schwartz, M. A. (2002) Zizimin1, a novel CDC42 activator, reveals a new GEF domain for RHO proteins. Nat Cell Biol 4, 639–647

4. Sakabe, I., Asai, A., Iijima, J., and Maruyama, M. (2012) Age-related guanine nucleotide exchange factor, mouse Zizimin2, induces filopodia in bone marrow-derived dendritic cells. Immunity & ageing : I & A 9, 2

5. Gadea, G., Sanz-Moreno, V., Self, A., Godi, A., and Marshall, C. J. (2008) DOCK10-mediated CDC42 activation is necessary for amoeboid invasion of melanoma cells. Curr Biol 18, 1456–1465

6. Nishikimi, A., Meller, N., Uekawa, N., Isobe, K., Schwartz, M. A., and Maruyama, M. (2005) Zizimin2: a novel, DOCK180-related CDC42 guanine nucleotide exchange factor expressed predominantly in lymphocytes. FEBS Lett 579, 1039–1046

7. Alcaraz-Garcia, M. J., Ruiz-Lafuente, N., Sebastian-Ruiz, S., Majado, M. J., Gonzalez-Garcia, C., Bernardo, M. V., Alvarez-Lopez, M. R., and Parrado, A. (2011) Human and mouse DOCK10 splicing isoforms with alternative first coding exon usage are differentially expressed in T and B lymphocytes. Human immunology 72, 531–537

8. Yelo, E., Bernardo, M. V., Gimeno, L., Alcaraz-Garcia, M. J., Majado, M. J., and Parrado, A. (2008) Dock10, a novel CZH protein selectively induced by interleukin-4 in human B lymphocytes. Molecular immunology 45, 3411–3418

9. Fluge, O., Bruland, O., Akslen, L. A., Lillehaug, J. R., and Varhaug, J. E. (2006) Gene expression in poorly differentiated papillary thyroid carcinomas. Thyroid 16, 161–175

10. Ruiz-Lafuente, N., Alcaraz-Garcia, M. J., Garcia-Serna, A. M., Sebastian-Ruiz, S., Moya-Quiles, M. R., Garcia-Alonso, A. M., and Parrado, A. (2015) Dock10, a CDC42 and RAC1 GEF, induces loss of elongation, filopodia, and ruffles in cervical cancer epithelial HeLa cells. Biology open 4, 627–635

11. Jaudon, F., Raynaud, F., Wehrle, R., Bellanger, J. M., Doulazmi, M., Vodjdani, G., Gasman, S., Fagni, L., Dusart, I., Debant, A., and Schmidt, S. (2015) The RHOGEF DOCK10 is essential for dendritic spine morphogenesis. Molecular biology of the cell 26, 2112–2127

12. Westcott, J. M., Prechtl, A. M., Maine, E. A., Dang, T. T., Esparza, M. A., Sun, H., Zhou, Y., Xie, Y., and Pearson, G. W. (2015) An epigenetically distinct breast cancer cell subpopulation promotes collective invasion. The Journal of clinical investigation 125, 1927–1943

13. Premkumar, L., Bobkov, A. A., Patel, M., Jaroszewski, L., Bankston, L. A., Stec, B., Vuori, K., Cote, J. F., and Liddington, R. C. (2010) Structural basis of membrane targeting by the Dock180 family of RHO family guanine exchange factors (Rho-GEFs). The Journal of biological chemistry 285, 13211–13222

14. Meller, N., Westbrook, M. J., Shannon, J. D., Guda, C., and Schwartz, M. A. (2008) Function of the N-terminus of zizimin1: autoinhibition and membrane targeting. Biochem J 409, 525–533

15. Yang, J., Zhang, Z., Roe, S. M., Marshall, C. J., and Barford, D. (2009) Activation of RHO GTPases by DOCK exchange factors is mediated by a nucleotide sensor. Science 325, 1398–1402 Mechanism of the dual specificity GEF DOCK10

16. Kulkarni, K., Yang, J., Zhang, Z., and Barford, D. (2011) Multiple Factors Confer Specific CDC42 and RAC Protein Activation by Dedicator of Cytokinesis (DOCK) Nucleotide Exchange Factors. The Journal of biological chemistry 286, 25341–25351

17. Harada, Y., Tanaka, Y., Terasawa, M., Pieczyk, M., Habiro, K., Katakai, T., Hanawa-Suetsugu, K., Kukimoto-Niino, M., Nishizaki, T., Shirouzu, M., Duan, X., Uruno, T., Nishikimi, A., Sanematsu, F., Yokoyama, S., Stein, J. V., Kinashi, T., and Fukui, Y. (2012) DOCK8 is a CDC42 activator critical for interstitial dendritic cell migration during immune responses. Blood 119, 4451–4461

18. Yamauchi, J., Miyamoto, Y., Chan, J. R., and Tanoue, A. (2008) ErbB2 directly activates the exchange factor Dock7 to promote Schwann cell migration. The Journal of cell biology 181, 351–365

19. Yamauchi, J., Miyamoto, Y., Hamasaki, H., Sanbe, A., Kusakawa, S., Nakamura, A., Tsumura, H., Maeda, M., Nemoto, N., Kawahara, K., Torii, T., and Tanoue, A. (2011) The atypical Guanine-nucleotide exchange factor, dock7, negatively regulates schwann cell differentiation and myelination. The Journal of neuroscience : the official journal of the Society for Neuroscience 31, 12579–12592

20. Kukimoto-Niino, M., Tsuda, K., Ihara, K., Mishima-Tsumagari, C., Honda, K., Ohsawa, N., and Shirouzu, M. (2019) Structural Basis for the Dual Substrate Specificity of DOCK7 Guanine Nucleotide Exchange Factor. Structure 27, 741–748 e743

21. Evans, P. (2006) Scaling and assessment of data quality. Acta Crystallogr D Biol Crystallogr 62, 72–82

22. Winn, M. D., Ballard, C. C., Cowtan, K. D., Dodson, E. J., Emsley, P., Evans, P. R., Keegan, R. M., Krissinel, E. B., Leslie, A. G., McCoy, A., McNicholas, S. J., Murshudov, G. N., Pannu, N. S., Potterton, E. A., Powell, H. R., Read, R. J., Vagin, A., and Wilson, K. S. (2011) Overview of the CCP4 suite and current developments. Acta Crystallogr D Biol Crystallogr 67, 235–242

23. McCoy, A. J., Grosse-Kunstleve, R.W., Adams, P.D.,Winn, M.D.,Storoni, L.C. and Read, R.J. (2007) Phaser crystallographic software. J. Appl. Cryst. 40, 658–674

24. Adams, P. D., Afonine, P. V., Bunkoczi, G., Chen, V. B., Davis, I. W., Echols, N., Headd, J. J., Hung, L. W., Kapral, G. J., Grosse-Kunstleve, R. W., McCoy, A. J., Moriarty, N. W., Oeffner, R., Read, R. J., Richardson, D. C., Richardson, J. S., Terwilliger, T. C., and Zwart, P. H. (2010) PHENIX: a comprehensive Python-based system for macromolecular structure solution. Acta Crystallogr D Biol Crystallogr 66, 213–221

25. Adams, P. D., Grosse-Kunstleve, R. W., Hung, L. W., Ioerger, T. R., McCoy, A. J., Moriarty, N. W., Read, R. J., Sacchettini, J. C., Sauter, N. K., and Terwilliger, T. C. (2002) PHENIX: building new software for automated crystallographic structure determination. Acta Crystallogr D Biol Crystallogr 58, 1948–1954

26. Schroder, G. F., Brunger, A. T., and Levitt, M. (2007) Combining efficient conformational sampling with a deformable elastic network model facilitates structure refinement at low resolution. Structure 15, 1630–1641

27. Davis, I. W., Leaver-Fay, A., Chen, V. B., Block, J. N., Kapral, G. J., Wang, X., Murray, L. W., Arendall, W. B., 3rd, Snoeyink, J., Richardson, J. S., and Richardson, D. C. (2007) MolProbity: all-atom contacts and structure validation for proteins and nucleic acids. Nucleic Acids Res 35, W375–383

28. Svergun, D., Barberato, C., and Koch, M. H. J. (1995) CRYSOL - a program to evaluate X-ray solution scattering of biological molecules from atomic coordinates Journal of Applied Crystallography 28, 768–773

29. Lewis, E. A., and Murphy, K. P. (2005) Isothermal titration calorimetry. Methods in molecular biology 305, 1–16

30. Velazquez-Campoy, A., Leavitt, S. A., and Freire, E. (2004) Characterization of protein-protein interactions by isothermal titration calorimetry. Methods in molecular biology 261, 35–54

31. Fisher, H. F., and Singh, N. (1995) Calorimetric methods for interpreting protein-ligand interactions. Methods in enzymology 259, 194–221

